# Anchoring functional connectivity to individual sulcal morphology yields insights in a pediatric study of reasoning

**DOI:** 10.1101/2024.04.18.590165

**Authors:** Suvi Häkkinen, Willa I. Voorhies, Ethan H. Willbrand, Yi-Heng Tsai, Thomas Gagnant, Jewelia K. Yao, Kevin S. Weiner, Silvia A. Bunge

## Abstract

A salient neuroanatomical feature of the human brain is its pronounced cortical folding, and there is mounting evidence that sulcal morphology is relevant to functional brain architecture and cognition. However, our understanding of the relationships between sulcal anatomy, brain activity, and behavior is still in its infancy. We previously found the depth of three small, shallow sulci in lateral prefrontal cortex (LPFC) was linked to reasoning performance in childhood and adolescence (Voorhies et al., 2021). These findings beg the question: what is the linking mechanism between sulcal morphology and cognition? To shed light on this question, we investigated functional connectivity among sulci in LPFC and lateral parietal cortex (LPC). We leveraged manual parcellations (21 sulci/hemisphere, total of 1806) and functional magnetic resonance (fMRI) data from a reasoning task from 43 participants aged 7–18 years (20 female). We conducted clustering and classification analyses of individual-level functional connectivity among sulci. Broadly, we found that 1) the connectivity patterns of individual sulci could be differentiated – and more accurately than rotated sulcal labels equated for size and shape; 2) sulcal connectivity did not consistently correspond with that of probabilistic labels or large-scale networks; 3) sulci clustered together into groups with similar patterns, not dictated by spatial proximity; and 4) across individuals, greater depth was associated with higher network centrality for several sulci under investigation. These results highlight that functional connectivity can be meaningfully anchored to individual sulcal anatomy, and demonstrate that functional network centrality can vary as a function of sulcal depth.

**Significance Statement:** A salient, and behaviorally relevant, feature of the human brain is its pronounced cortical folding. However, the links between sulcal anatomy and brain function are still poorly understood – particularly for small, shallow, individually variable sulci in association cortices. Here, focusing on individually defined sulci in lateral prefrontal and parietal regions, we offer a novel, anatomically informed approach to defining functional connectomes. Further, we demonstrate, for the first time, a link between functional network centrality and sulcal morphology.

## Introduction

A salient feature of the human brain is its pronounced cortical folding. Indeed, 60–70% of the cortex is buried in sulci (Welker, 1990; Armstrong et al., 1995; Zilles et al., 2013). Sulci develop prenatally under varying degrees of genetic control through a combination of biomechanical forces, including white matter tract formation, and differential cortical tissue outgrowth (Im and Grant, 2019; Van Essen et al., 2007; Zilles et al., 2013). Most sulci can be identified in every brain; however, they vary across individuals in location and shape –– especially those within association cortices that have expanded the most throughout evolution, referred to here as putative tertiary sulci (pTS).

There is mounting evidence that individual variability in sulcal morphology is functionally and behaviorally relevant across age groups, species, and clinical populations (Artiges et al., 2006; Bouhali et al., 2024; Cachia et al., 2018; Fedeli et al., 2022; Garrison et al., 2015; Miller and Weiner, 2022; Maboudian et al., 2024; Natu et al., 2021). Individual variability in sulcal spatial configuration and morphology was hypothesized decades ago to impact neural efficiency and cognition (Sanides, 1962, 1964). Consistent with this hypothesis, features of specific sulci have been linked to variability in a variety of cognitive tasks (Aichelburg et al., 2016; Borst et al., 2014; McGugin et al., 2020; Meredith et al., 2012; Santacroce et al., 2024; Li et al., 2024; Tissier et al., 2018; Parker et al., 2023; Voorhies et al., 2021; Willbrand et al., 2023a, 2023b, 2023c; Yao et al., 2023).

In parallel to research linking sulcal morphology to cognition, ongoing work links sulcal morphology to functional brain organization (Amiez and Petrides, 2014; Amiez et al., 2013; Bodin et al., 2018; Cordeau et al., 2023; Derrfuss et al., 2009; Eichert et al., 2021; Germann et al., 2020; Huster et al., 2014; Sun et al., 2016; Weiner et al., 2014; Zlatkina et al., 2016), including network connectivity. For example, the presence or absence of a specific sulcus in one part of the brain can affect functional connectivity in other lobes (Lopez-Persem et al., 2019). Further, adjacent pTS can participate in different large-scale networks, leading to the proposal that pTS be used as an individual coordinate space for functional connectomes, adopting one facet of precision imaging (Miller et al., 2021; Miller and Weiner, 2022).

Prior work implementing a model-based approach identified three pTS in LPFC whose depth was related to abstract reasoning task performance across a sample of children and adolescents (Voorhies et al., 2021). Thus, a major impetus for this study was to examine the functional relevance of sulcal depth: specifically, a possible association with network centrality. We focused on sulci in LPFC and lateral parietal cortex (LPC): areas that work together to support reasoning and other higher cognitive functions (e.g., Vendetti and Bunge, 2014; Krawczyk et al., 2011; Stuss and Knight, 2013; Woolgar et al., 2010). In particular, a study including the present pediatric sample showed that strength of LPFC-LPC white matter and functional connectivity were linked to reasoning performance (Wendelken et al., 2017).

To test for depth-connectivity relations, we first characterized functional connectivity patterns among sulci – i.e., a *sulcal functional connectome.* To do so, we leveraged manual labels of 1806 sulci (both large and pTS in LPFC and LPC, including newly identified LPC sulci (Willbrand et al., 2023c) from 43 participants aged 7–18 years. We then followed a four-pronged approach. First, we tested whether manually labeled sulci could be distinguished based on their patterns of functional connectivity. Second, we tested how sulci clustered based on connectivity patterns. Third, we computed graph metrics of network centrality and tested whether greater depth of the three pTS previously implicated in reasoning was associated with higher centrality. Fourth, we conduced control analyses to validate our novel sulcal connectomics approach. Altogether, we propose that anchoring functional connectivity to sulcal anatomy is a promising way forward, and demonstrate a novel link between sulcal anatomy and network centrality.

## Materials and Methods

### Participants

Participants were selected from the Neurodevelopment of Reasoning Ability (NORA) dataset (Wendelken et al., 2011). Participants were all right-handed native English speakers, ranging in age from 7 to 18 (**Table 1**). 25/43 of these participants were also included in the prior morphological study (Voorhies et al., 2021). All participants underwent preliminary screening as part of the NORA study to include only right-handed, neurotypical children and adolescents. Participants and their parents gave their informed assent and/or consent to participate in the study, which was approved by the Committee for the Protection of Human Participants at the University of California, Berkeley.

**Table 1.** Participant characteristics and demographics.

|  |  |
| --- | --- |
| Age (years), mean (range) | 14.26 (7.12–18.86) |
| Gender (female/male) | 20/23 |
| Matrix reasoning (t-score), mean (range) | 58.43 (43.00–81.00) |
| Head motion (mean framewise displacement in mm), mean (range) | 0.22 (0.08–0.43) |
| fMRI data after censoring (minutes), mean (range) | 12.67 (5.47–16.20) |
| Annual Household Income (32 responses) |  |
| \$16,000 to \$99,999 | 14 |
| \$100,000 to \$200,000 | 15 |
| Over \$200,000 | 3 |
| Highest Degree Earned by Parent/Guardian (36 responses) |  |
| High School/GED/Associate degree | 9 |
| Bachelor's degree | 13 |
| Master's degree/Doctorate/Professional | 14 |
| Racial Categories (41 responses) |  |
| White | 20 |
| Black or African American | 7 |
| Asian | 2 |
| More Than One Race | 12 |

Parent/guardian-reported demographics. Only ~50% of participants responded to the question of whether they were Hispanic or Latino. The Matrix Reasoning score was not available for one participant.

Participants were selected from the NORA study based on screening for usable T1 and fMRI data on any assessment, using MRIQC (23.1.0, Esteban et al., 2017) and manual inspection for scanner artifacts using FSLeyes (McCarthy, 2018). For fMRI, we required three complete runs of fMRI with 165 volumes and estimated gross motion measured by mean framewise displacement < 0.5 mm (Power et al., 2014), and runs combined > 5 min worth of frames free from significant motion (framewise displacement < 0.5 mm or DVARS < 1.5). These criteria led us to retain 46% (49/107) of participants. Of these, two were excluded because their LPFC sulci had not been labeled. Four additional participants were excluded because they were missing one of the pTS that is not present universally (one participant did not have a rh.pimfs in LPFC, and three participants did not have a lh.slocs-v in LPC); this reflects normal variation for these two sulci. Thus, 43 participants with all sulci present in both hemispheres were included in our analyses.

The age and gender distributions of these participants are reported in **Table 1**, along with their performance on a standard test of reasoning (Wechsler Intelligence Scale for Children (WISC-IV) Matrix Reasoning task; Wechsler, 1949), degree of head motion, and amount of high-quality fMRI data used in analyses.

Parent/guardian-reported demographics. Only ∼50% of participants responded to the question of whether they were Hispanic or Latino. The Matrix Reasoning score was not available for one participant.

### fMRI task

Functional connectivity was measured based on fMRI data collected during performance of a reasoning task (**Figure 1**; see also Wendelken et al., 2011). The data were derived from a blocked fMRI task design, with three runs of 5 minutes 25 seconds each, for a total of 16.3 minutes. The current analyses focus on functional connectivity across all task and rest blocks; thus, we measured general functional connectivity (Elliott et al., 2019) in the context of a reasoning task.

**Figure 1.**
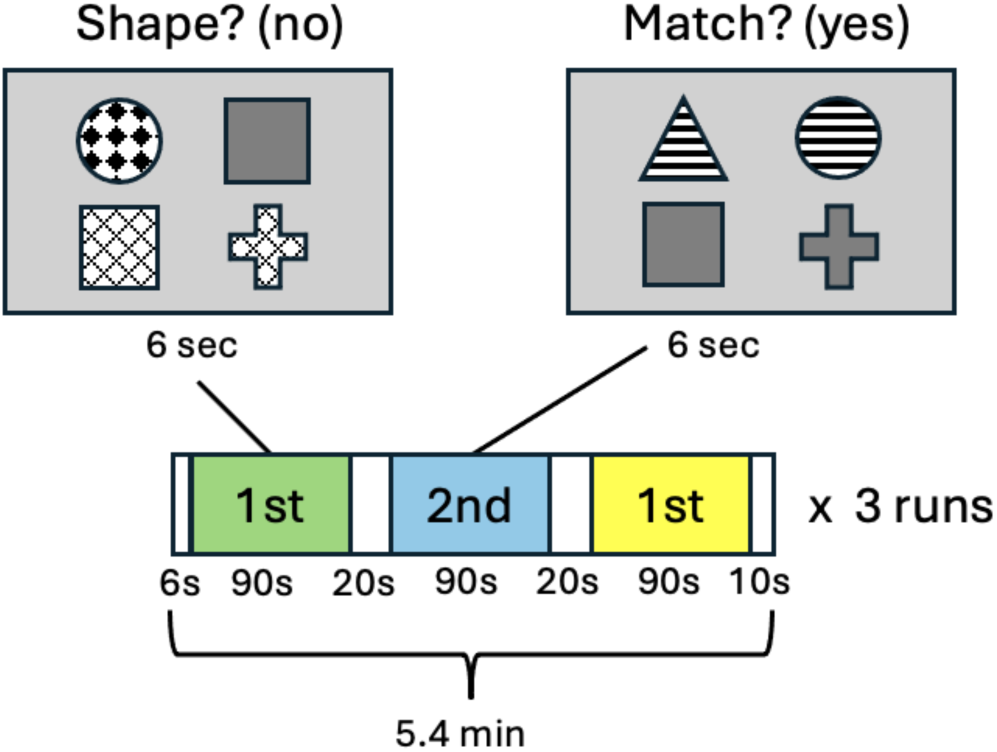
fMRI task paradigm. Each participant completed three runs of 5.4 min each of a reasoning task. Each run included three 90-s task blocks of 15 trials of 6 seconds each: two 1st-order blocks – one shape block and one pattern block (depicted in green and yellow, respectively) – and one 2^nd^-order block (blue). Task blocks alternated between 1st- and 2nd- order within each run, separated by 20-s rests, along with a 6-s and 10-s rest at the start and end of each run, respectively. On 1st-order trials, participants saw an array of four shapes with various pattern fills and indicated whether either row of stimuli had the same shape (the Shape condition) or fill pattern (the Pattern condition). In the shape trial depicted, neither row of stimuli have the same shape; thus, the correct answer is ‘no’. On 2nd-order (Match) trials, participants indicated whether the two rows of stimuli in the array matched on the same feature (shape or pattern). In the match trial depicted, both rows of stimuli match on pattern; thus, the correct answer is ‘yes’.

Each run included three 90-second task blocks, one for each of three task conditions described below **(Figure 1**). The task blocks were separated by 20-seconds of rest, with 6- and 10-second rest at the start and end of each scan, respectively. In total, there were 13.5 minutes of task and 2.8 minutes of rest across the scan session. All participants performed the task well above chance-level accuracy, as reported in a previous fMRI study involving 6–18- year-olds from which the present sample was drawn (mean ± SD for 1st-order relations: 92.8 ± 6.6%; 2nd-order relations: 90.0 ± 9.6%; Wendelken et al., 2011).

With regard to the specific cognitive demands of the task, participants viewed an array of four simple patterned shapes and had to make a decision about the relations between two stimuli in a row (1st-order relations) or the relation between the two rows of stimuli (2nd-order relations). There were three task conditions, presented in separate blocks of 15 trials each: 1st-order shape relations, 1st-order pattern relations, and 2nd-order relations (for a total of 135 trials). In the 1st-order conditions, participants judged whether the pair of stimuli in each row shared a particular feature: shape or fill pattern (e.g., on a shape trial, judging that a checkered circle and a solid square do not match on shape). In second-order reasoning blocks, they judged whether two pairs of stimuli matched according to the same feature – that is, the shape or fill pattern (e.g., judging that the top pair (a striped triangle and circle) and bottom pair (a solid square and cross) both match according to pattern).

### MRI data acquisition

Brain imaging data were collected on a Siemens 3T Trio system at the University of California Berkeley Brain Imaging Center. T1-weighted MPRAGE anatomical scans (TR = 2300 ms, TE = 2.98 ms, 1 × 1 × 1 mm voxels) were acquired for cortical morphometric analyses. fMRI data were acquired using gradient-echo EPI sequence, TR = 2000 ms, TE = 25 ms, 33 axial slices, 2.0 × 1.8 × 3.0 mm voxels, no interslice gap, flip angle = 90°, field of view = 230 mm, 120 volumes per run).

### Cortical surface reconstruction

FreeSurfer’s automated segmentation tools (Dale et al., 1999; Fischl and Dale, 2000) (FreeSurfer 7.1.0) were used to generate cortical surface reconstructions. Each anatomical T1-weighted image was segmented to separate gray from white matter, and the resulting boundary was used to reconstruct the cortical surface for each participant (Dale et al., 1999; Wandell et al., 2000). Each reconstruction was visually inspected for segmentation errors, and these were manually corrected when necessary.

### Manual labeling of LPFC and LPC sulci

To investigate the network defined by the LPFC–LPC sulcal features, 42 sulci (10 LPFC and 11 LPC in each hemisphere) were manually defined for each participant and hemisphere.

These anatomical features were defined according to the most recent atlas by Petrides (Petrides, 2019) and our previous work (Voorhies et al., 2021; Willbrand et al., 2023c). The anatomical locations are illustrated in **Figure 2**, and all 1806 sulcal definitions in all participants can be found in **Extended Data** Figure 2-1. The location of each sulcus was confirmed by trained independent raters (W.I.V., E.H.W., J.Y., T.G., Y.T.) and finalized by a neuroanatomist (K.S.W.). Surface vertices for each sulcus were selected using tools in FreeSurfer, guided by the pial and smoothwm surfaces (Weiner et al., 2018; Miller et al., 2021), and saved as surface labels for vertex-level analysis. We have indicated pTS with asterisks in Figures 2, 4–7; however, future studies using novel in-vivo developmental imaging in relation to classic post-mortem techniques are required to definitively determine which are secondary vs. tertiary sulci based on their emergence in gestation (Chi et al., 1977) in addition to their depth and surface area.

**Figure 2.**
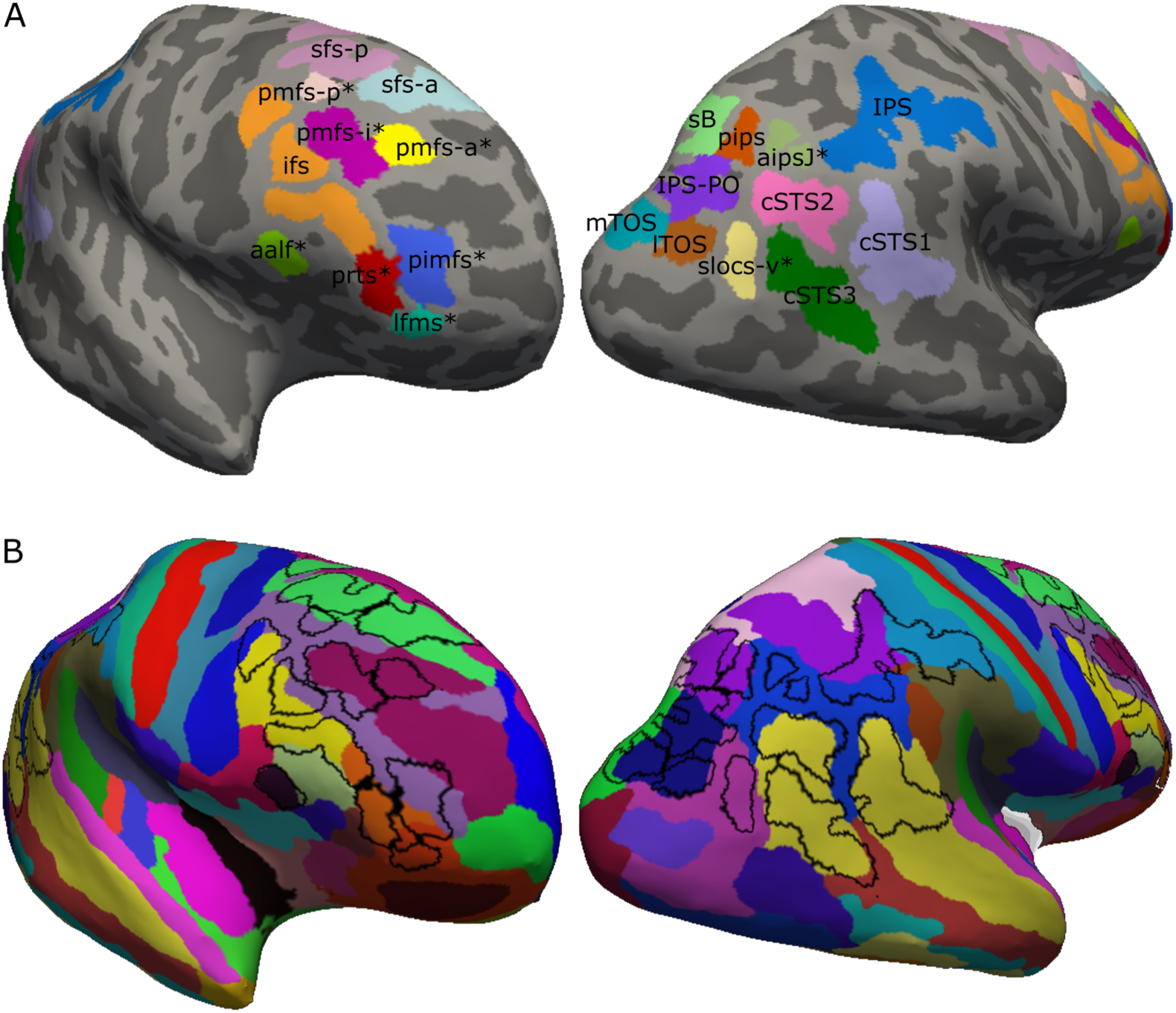
Lateral prefrontal (LPFC) and lateral parietal (LPC) sulcal definitions. (A) Manual sulcal definitions on an example cortical surface reconstruction of a right hemisphere. *Putative tertiary sulcus (pTS). See **Extended Data** Figure 2-1 for individual sulcal definitions for all participants. (B) Manual definitions (black outlines) overlaid on the Destrieux atlas generated using automated methods (Fischl, 2004; Destrieux et al., 2010). Note that many of the LPFC and LPC sulci (black outlines) do not align with the automated definitions of these structures. Abbreviations: ifs: inferior frontal sulcus; sfs-p: superior frontal sulcus-posterior; sfs-a: superior frontal sulcus-anterior; pmfs-p: posterior middle frontal sulcus-posterior; pmfs-i: posterior middle frontal sulcus-intermediate; pmfs-a: posterior middle frontal sulcus-anterior; pimfs (combining pimfs-v and pimfs-d): para-intermediate frontal sulcus; aalf: ascending ramus of the lateral fissure; prts: pretriangular sulcus; lfms: lateral frontomarginal sulcus; sB: sulcus of Brissaud; pips: intermediate parietal sulcus-posterior; mTOS: transverse occipital sulci-medial; lTOS: transverse occipital sulcus- lateral; IPS-PO: paroccipital intraparietal sulcus; IPS: intraparietal sulcus; cSTS1,2,3: three branches of the caudal superior temporal sulcus; aipsJ: intermediate parietal sulcus of Jensen- anterior; slocs-v: supralateral occipital sulcus-ventral.

### Morphological features

Sulcal depth was defined by maximum ‘sulc’ value estimated by FreeSurfer for each sulcal label. Sulcal area was defined by total surface area (mris_anatomical_stats).

### fMRI data preprocessing

We drew the raw fMRI data for included participants from the previous dataset, and preprocessed them de novo. Here, we used fMRIprep version 21.0.1 (Esteban et al., 2019)[RRID:SCR_016216], which is based on Nipype 1.6.1 (Gorgolewski et al., 2011)[RRID:SCR_002502]. Each T1w (T1-weighted) volume was corrected for intensity non-uniformity using N4BiasFieldCorrection v2.3.3 (Tustison et al., 2010) and skull-stripped using antsBrainExtraction.sh v2.1.0 (using the OASIS template). Brain surfaces were reconstructed using recon-all from FreeSurfer v7.1.0 (Dale et al., 1999)[RRID:SCR_001847], and the brain mask estimated previously was refined with a custom variation of the method to reconcile ANTs-derived and FreeSurfer-derived segmentations of the cortical gray-matter of Mindboggle (Klein et al., 2017)[RRID:SCR_002438]. Brain tissue segmentation of cerebrospinal fluid (CSF), white matter (WM) and gray matter (GM) was performed on the brain-extracted T1w using fast (Zhang et al., 2001, FSL v6.0.5, RRID:SCR_002823).

For each of the three runs, the initial three volumes were excluded, functional data was slice time corrected using 3dTshift from AFNI v16.2.07 (Cox, 1996)[RRID:SCR_005927] and motion corrected using mcflirt (Jenkinson et al., 2002)(FSL v6.0.5). This was followed by co-registration to the corresponding T1w using boundary-based registration (Greve and Fischl, 2009) with six degrees of freedom, using bbregister (FreeSurfer v7.1.0). Motion correcting transformations, BOLD-to-T1w transformation and T1w-to-template (MNI) warp were concatenated and applied in a single step using antsApplyTransforms (ANTs v2.1.0) using Lanczos interpolation. The functional series were then sampled to native mid-grey matter surface by averaging across the cortical ribbon.

Physiological noise regressors were extracted by applying CompCor (Behzadi et al., 2007). For aCompCor, six principal components were calculated within the intersection of the subcortical mask and the union of CSF and WM masks calculated in T1w space, after their projection to the native space of each functional run. Framewise displacement (Power et al., 2014) was calculated for each functional run using the implementation of Nipype.

Following preprocessing via fMRIPrep, fMRI data were despiked (AFNI 3Ddespike, default c1 = 2.5, c2 = 4.0) and bandpass filtered (AFNI 3DBandpass, 0.008–0.1 Hz).

Nuisance regressors were filtered and regressed from the timeseries (Nilearn image.clean_img). The confounds included linear trends, the first 5 aCompCor components, and 24 motion parameters (Friston et al., 1996). In addition to the tools specifically mentioned, this denoising and further data analysis relied on open-source Python packages including NiBabel [RRID:SCR_002498](Brett et al., 2022), NumPy [RRID:SCR_008633](Harris et al., 2020), SciPy [RRID:SCR_008058](Virtanen et al., 2020) and Pandas [RRID:RRID:SCR_018214](McKinney, 2010).

### Measurement of sulcal functional connectivity and correction for spatial autocorrelations

We computed functional connectivity between all 42 sulcal labels for each participant. To this end, the mean timeseries of all vertices in each sulcal label were averaged. Functional connectivity was measured by pairwise Pearson correlations between the timeseries for each sulcal label across all three runs, scrubbing frames associated with large motion spikes (framewise displacement < 0.5 or DVARS < 1.5). The correlation values were subsequently normalized to z-scores via a Fisher transformation, and then combined into a 42 × 42 inter-regional correlation matrix for each participant. The analyses were based on the full acquisition with task and rest blocks concatenated. To further mitigate effects of in-scanner head motion that might skew network definition for example by inflating short-distance connectivity, mean framewise displacement was residualized from the data prior to analyses of network organization and included as a covariate in analyses on associations with age.

The results featured here control for spatial autocorrelation, which may artificially inflate functional connectivity between adjacent sulcal components. Spatial autocorrelation was estimated and controlled for per participant and hemisphere as follows: 1) Functional connectivity and corresponding geodesic distances were calculated for nodes in 32k meshes (Connectome Workbench, v1.4.2, RRID:SCR_008750, Marcus et al., 2011) of cortical mid-grey matter, excluding the medial wall (Python library tvb-gdist 1.0.3; https://github.com/the-virtual-brain/tvb-gdist). 2) Spatial autocorrelation was modeled as a function defined by a rate of exponential distance-dependent decrease and the constant baseline level to which it decays, and fitted to the data divided into 1 mm bins using gradient descent, similar to Shinn et al. (2023). For all participants and hemispheres, the positive bias of adjacent connections was found to taper off at < 1 cm. To define a regressor that describes the short-distance positive bias and does not change the connectivity values of sulcal pairs at longer distances than the participant’s estimated range of spatial autocorrelations, the constant baseline level was subtracted from the fitted function. 3) Minimum geodesic distance was extracted for the vertices of each sulcal label pair of the participant. 4) Based on the autocorrelation function and calculated minimum distance, each sulcal label pair was matched with the expected positive bias. 5) Association between the bias regressor and connectivity of the sulcal network was estimated for each hemisphere and participant using least squares regression.

Connectivity values were then residualized for the estimated regression with the bias variable. 6) The corrected within-hemisphere connections were combined with between-hemisphere connections for further analysis.

While spatial autocorrelations are an important potential confound, correcting for them may diminish true short-distance connectivity and confound network topological measures (Shinn et al., 2023). Thus, we also ran supplementary analyses in which we did not correct for them.

### Discriminability of sulcal connectivity patterns

We performed a classification analysis testing the discriminability of the sulci from one another based on their connectivity patterns. The multiclass classification problem was split into 861 binary classification problems (one-vs-one approach) (Galar et al., 2011). We chose a support vector machine (SVM) as a classifier, given the small, high-dimensional sample, and employed a linear kernel to avoid overfitting. SVM were implemented using scikit-learn [RRID:SCR_002577](Pedregosa et al., 2011) (based on LIBSVM, default regularization C = 1), and trained to discriminate each pair of sulci based on their functional connectivity fingerprint. To avoid having the classifications be driven by strong autocorrelations, the connectivity between the two sulci being tested was excluded from each binary classification, such that the classification was based on the connectivity strength between each of these two sulci and the 40 remaining sulci. Connectivity weights of each participant were normalized to [0,1], and each feature to [0,1] during cross-validation. Classification performance was assessed using leave 3 participants out cross-validation repeated 10 times, and accuracy (the percentage of samples predicted correctly) was measured as the mean value across folds.

Empirical p-values were generated against the null hypothesis that connectivity fingerprints and sulcal labels are independent, using permutation testing (100 permutations of each fold) and correcting for multiple comparisons using the Benjamini-Hochberg false discovery rate (FDR) (Benjamini and Hochberg, 1995).

### Control analyses evaluating sulcal functional specificity

To test whether sulci are functionally relevant, we conducted control analyses comparing functional connectivity and classification accuracy for manually labeled sulci vs. null models generated by spin testing (Alexander-Bloch et al., 2018). For each sulcal definition in each individual, we generated 1000 rotated parcels of the same size and shape (code adapted from https://github.com/netneurolab/markello_spatialnulls; Markello and Misik, 2021). These rotated sulcal definitions were required to fall within the envelope of the original LPFC or LPC sulci and were combined into 1000 rotated sulcal networks per participant (**Figure 3A**). We then compared functional connectivity strength and classification accuracy between the manual and rotated labels.

**Figure 3.**
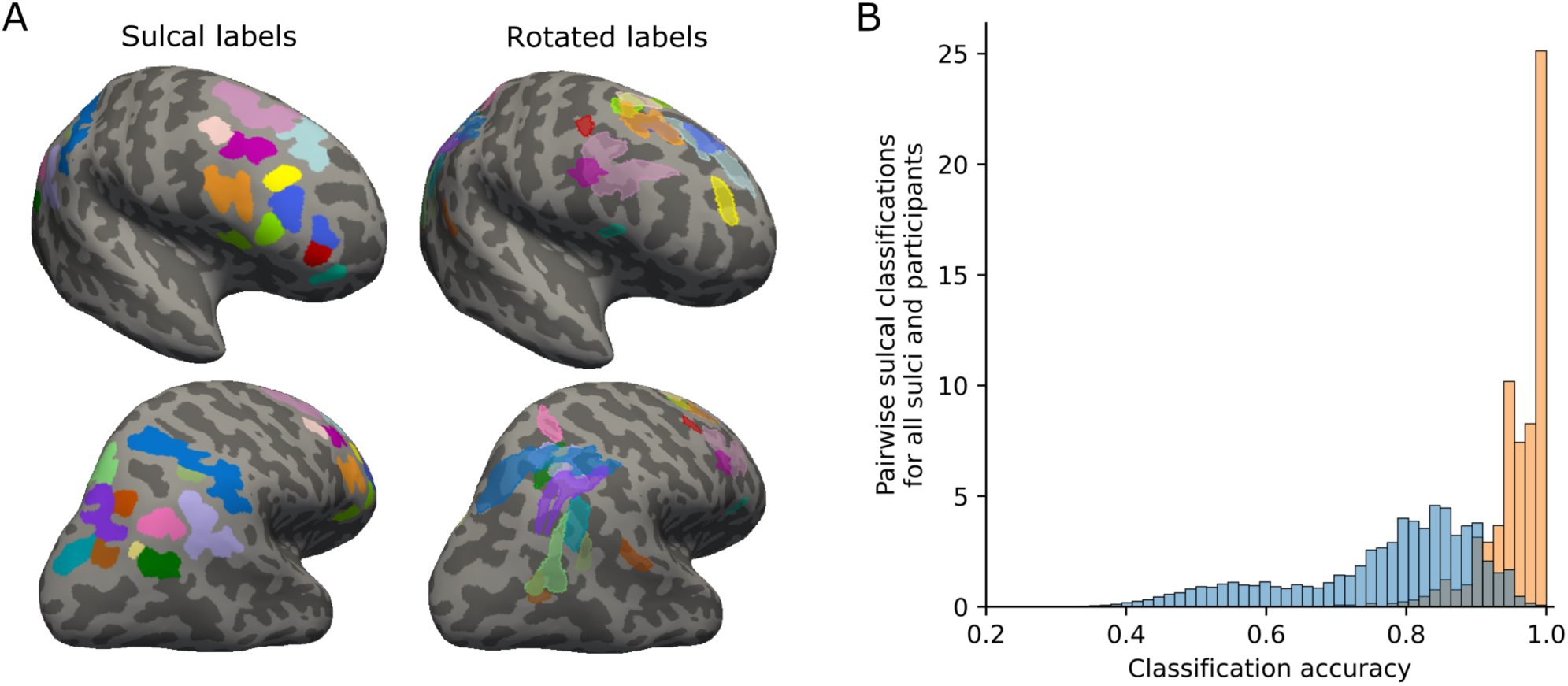
**Sulcal specificity of functional connectivity fingerprints compared to null models generated by spin testing**. (A) Manual sulcal definitions in LPFC (upper row) and LPC (lower row) and a set of rotated labels in one of 1000 permuted networks, in an example participant. Colors match Figure 2A. (B) Density plot of classification accuracy for selected manual sulcal definitions and the corresponding null model (1000 rotated labels per sulcus), showing the number of accurate classifications across all sulcal pairs and participants. Classification accuracy of 1 would correspond to perfect discrimination for a sulcal pair. The plot shows results for the target sulci in LPFC (bilateral pimfs, pmfs-a, pmfs-i, pmfs-p), as well as the pTS in LPC included in exploratory analyses (aipsJ, slocs-v).

### Probabilistic sulcal labels

To test the necessity of manual labeling in individual participants – a laborious process that requires anatomical expertise – we computed probability maps for all sulci based on cortical overlap between participants. Probabilistic labels were generated for each participant based on the remaining (N-1) participants in the cohort, establishing spatial correspondence between participants using spherical surface registration to the fsaverage template (Fischl et al., 1999). Probabilistic labels were thresholded to include vertices that were included in a given sulcal label for at least 33% of participants. In cases of overlap between probabilistic labels, we assigned a vertex to the sulcus with greater overlap between participants.

Despite this liberal clustering threshold, four highly variable sulci did not have any vertices in their probabilistic definition (lh.slocs-v and rh.aipsJ in all 43 participants, lh.aipsJ and rh.pmfs_a in 42); this, in and of itself, speaks to the fact that probabilistic labels (not to mention standard atlases) do not always accurately represent the anatomy of individual participants. Sulcal networks based on the existing probabilistic labels were then calculated as in the main analysis.

### Connectivity-based clustering

To detect groups of sulci with similar sulcal connectivity profiles, connectivity matrices were averaged into a mean connectivity matrix, and the mean matrix was clustered using an unsupervised community detection algorithm (Infomap) (Rosvall and Bergstrom, 2008).

Clusters were investigated at density thresholds 0.01–0.5 in steps of 0.01, preserving the indicated proportion of the strongest weights at each iteration. Results at different thresholds were combined into a “consensus” clustering (similar to e.g. Gordon et al., 2017; Hwang et al., 2017; Laumann et al., 2015; Marek et al., 2019; Moore et al., 2024; Power et al., 2011; Rajesh et al., 2024), i.e., we did not consider the sparsity of the network. This grouping should be considered a rough summary view mainly used for visualization, also given that the group-level clusters may not mirror individual clustering (Laumann et al., 2015; Smith et al., 2023).

The representativeness of the group-level clustering at the individual participant level was assessed as follows. First, clustering was applied to individual-level networks. For each label pair, co-clustering was defined as the number of times each pair of labels was found in the same cluster, at clustering thresholds ranging from the lowest threshold at which the sulcus was clustered with at least one other sulcus to the threshold at which all sulci were clustered into one. Co-clustering counts were collapsed into a binary estimate indicating whether the pair clustered together for at least three of the investigated thresholds. Finally, an agreement matrix described the frequency of co-clustering per sulcal pair, presented as a proportion of participants.

### Sulcal associations with canonical large-scale resting-state fMRI networks

To determine how individual sulci associate with previously characterized large-scale functional networks, we assessed the similarity of each sulcus and 14 networks in the Masonic Institute for the Developing Brain (MIDB) Precision Brain Atlas (Hermosillo et al., 2024), derived from probabilistic network maps from resting-state fMRI data, that were largely invariant in network topography across roughly 6000 9–10 year-olds. Network seed masks were defined at 61% probability in the atlas, producing reasonably sized seeds for connectivity analysis. The atlas labels were projected from standard space (fs_LR_32) to the native surfaces of each participant using Connectome Workbench tools and custom scripts. The mean timeseries of each network label and sulcus (same hemisphere) were then correlated as above.

### Graph network metrics

Based on the 42 × 42 matrices of sulcal connectivity, we derived graph metrics for each node (sulcus) in each participant. Individual nodes were characterized using three topological measures describing their level of centrality in the LPFC-LPC sulcal network: 1) degree, or the number of links connected to the node; 2) betweenness, or the frequency that a node is on the shortest path connecting other nodes; and 3) participation coefficient, a measure of how evenly distributed a node’s connections are across clusters (Rubinov and Sporns, 2010). To account for differences in overall connectivity strength across participants, the graph analyses used binarized networks and area under the curve (AUC) estimates, integrating values across 0.01–0.2 density. Participation coefficient was calculated using the group-level clusters shown in **Figure 4B**.

**Figure 4.**
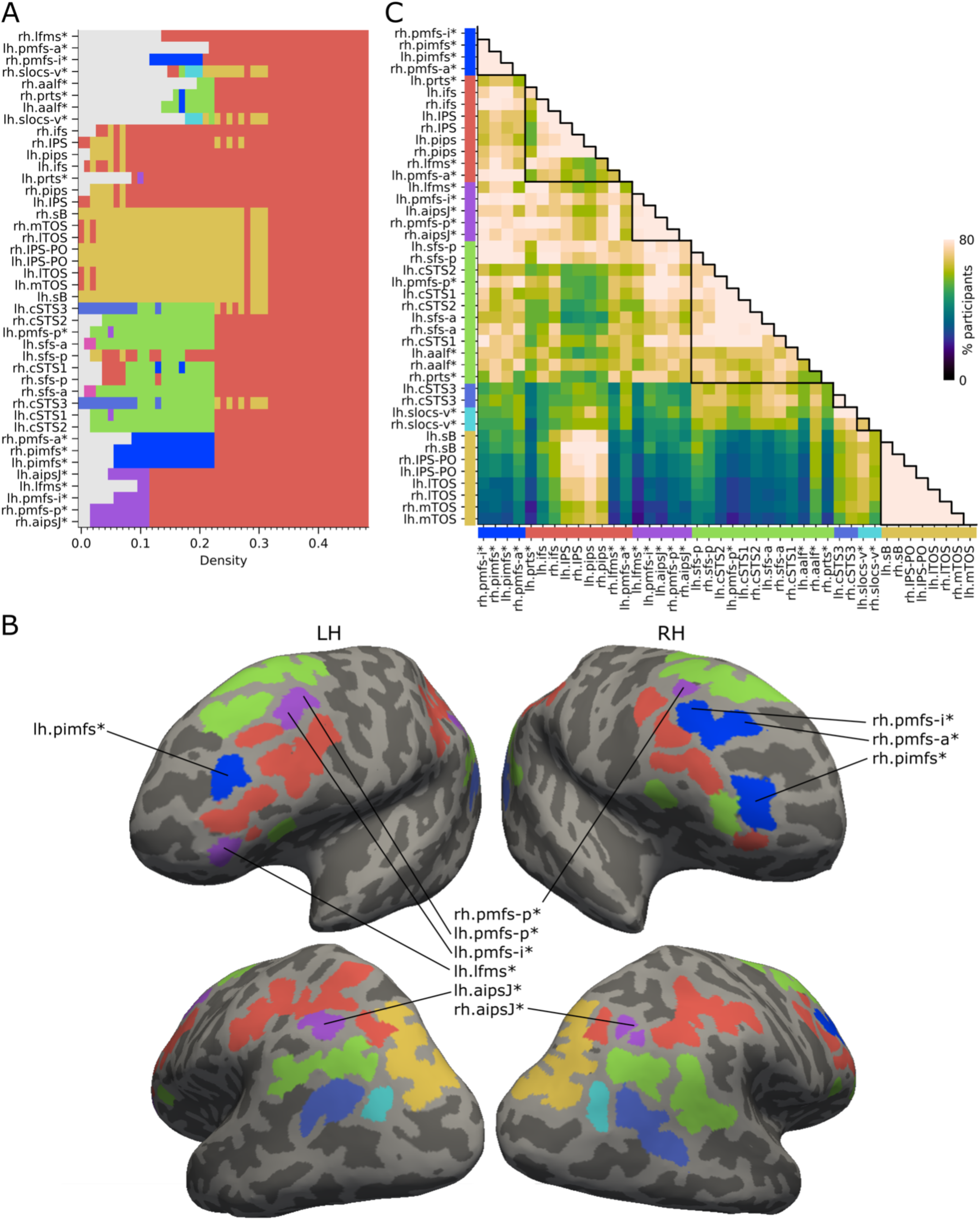
Lateral frontoparietal functional connectivity based on individual sulcal morphology. (A) Group-averaged (N = 43) network clusters identified at 0.01–0.5 density thresholds. Colors reflect cluster identity; white refers to nodes that are isolated or form a single-sulcus cluster at the threshold. *Putative tertiary sulcus (pTS). (B) Clusters selected to capture distinctions across at least two thresholds in the group results, displayed on the cortical surfaces of an example participant. The colors correspond to those in panel A. (C) Co-clustering at the individual level in relation to the group-level results. This matrix displays the percentage of participants for whom two sulci clustered together in individual-level clustering analyses. The sulci are grouped based on the group-level clustering results, as indicated by the colored bars along the axes. Lighter colors indicate sulcal pairs that clustered together at more than one threshold for a larger number of participants. Pairs of sulci assigned to the same group-level cluster are delineated by black outlines; relatively strong agreement across participants is observed for these pairs.

For each graph metric, the sulcal components scoring consistently high across participants were defined by comparison to the mean of all values for a given participant using two-tailed paired t-tests, and FDR-corrected for multiple comparisons across all 42 nodes.

Graph metrics were computed using the Python port (https://pypi.org/project/bctpy) of Brain Connectivity Toolbox (BCT; Rubinov and Sporns, 2010). Statistics were computed using Scipy and statsmodels[RRID:SCR_016074](Seabold and Perktold, 2010). For matrix visualization, nodes were ordered both by clusters and within each cluster to maximize the number of edges close to the main diagonal, using simulated annealing (BCT reorder_matrix). Visualizations were created using Matplotlib [RRID:SCR_008624](Hunter, 2007), Seaborn[RRID:SCR_018132](Waskom, 2021) and NetworkX [RRID:SCR_016864](Hagberg et al., 2008). Spring layout was generated using the Kamada-Kawai method (Kamada and Kawai, 1989).

### Testing associations between sulcal depth and network centrality for three pTS in LPFC

Building on the previous finding that the depth of right pimfs, pmfs-a, and pmfs-i pTS (all assigned to the blue cluster, **Figure 4B**) predicted reasoning performance beyond age (Voorhies et al., 2021), we tested whether the depth of these sulci also related to differences in functional organization during reasoning. First, we tested, for the left and right hemisphere counterpart of each sulcus, whether its depth relates to its graph metrics (nodal degree, betweenness, and/or participation coefficient). Associations between depth and each metric were tested using ordinary least squares regression, controlling for age effects on depth and functional connectivity by including it as an additional explanatory variable (two-tailed tests, FDR-corrected across the 18 combinations of the six sulci and three graph metrics). We followed up with analogous regression analyses with individual edges as dependent variables to pinpoint the pairwise connections that drove the significant associations (one-tailed tests, FDR-corrected across the 42 connections investigated per sulcus).

### Code accessibility

Scripts to perform data preprocessing and statistical analyses will be freely available with the publication of the paper on GitHub (https://github.com/cnl-berkeley/stable_projects).

Requests for further information or raw data should be directed to the corresponding authors.

## Results

### Classification analyses

#### Sulci exhibited dissociable functional connectivity profiles

To assess whether (or which) sulci could be differentiated on the basis of their functional connectivity, we tested whether any two sulci for a given participant could be classified based on their connectivity fingerprints. Indeed, the SVM classifier could successfully discriminate each pair of sulci well above chance level (i.e., 50%) across participants, with very high accuracy for most – but not all – pairs of sulci (mean: 96%, range: 71–100%; FDR-corrected p < 0.01). Overall, then, sulci generally had differentiable connectivity profiles, whereby two sulci could be distinguished on the basis of their functional connectivity with 96% accuracy on average.

#### Functional connectivity was meaningfully related to sulcal anatomy

To test whether sulci are functionally relevant, we conducted control analyses comparing functional connectivity and classification accuracy for manually labeled sulci vs. null models generated by spin testing (Alexander-Bloch et al., 2018), involving 1000 rotated parcels of the same size and shape for each manually labeled sulcus.

We first examined overall distributions of functional connectivity among all pairs of sulcal vs. rotated labels. We observed a more dynamic range of correlation values for actual sulci than for rotated labels – including both stronger and weaker values (sulci: mean 0.29, range -0.52–1.24, SD 0.27; rotated labels: mean 0.27, range -0.13–0.91, SD 0.14). Levene’s test confirmed that the variances for pairwise connectivity strength were not equal, F1,5674 = 857.70, p < 0.0001). Critically, we then tested whether pairwise classification accuracy across participants was higher for veridical sulcal networks than our null model. As predicted, we confirmed that classification accuracy was significantly higher for sulcal labels than for rotated labels (**Figure 3B**; mean accuracy [range] for the three target sulci bilaterally: 0.93 [0.78–1.0]; rotated labels: 0.74 [0.50–.87]; t164 = 34.69; p < 0.0001). Thus, compared with equally sized and shaped random patches of cortex, actual sulci had a greater dynamic range of connectivity values, indicating a more sensitive unit of analysis, and the pTS under investigation had substantially more discriminable connectivity profiles.

### Clustering analyses

#### Sulcal clustering within and across LPFC and LPC on the basis of functional connectivity

Next, we sought to test whether sulci clustered into groups with similar connectivity profiles. We first conducted this analysis at the group level; we then conducted a co-clustering analysis at the individual level to quantify the extent to which these clusters represented individual-level connectivity.

We identified five sulcal groupings in the 0.01–0.5 density range at the group level (**Figure 4A**): one consisting of LPFC pTS (dark blue), one of pTS in both LPFC and LPC (purple), and two consisting of large sulci and pTS in both LPFC and LPC (red, green), and one of larger LPC sulci (yellow). In addition, two sulci (slocs-v, cyan; cSTS3, medium blue) formed two distinct clusters with their interhemispheric counterparts. This analysis reveals, for the first time, sulcal-functional coupling both within and between LPFC and LPC sulci (**Figure 4B**).

Focusing on these five clusters derived from the group mean network (**Figure 4B**), we next assessed whether they generalized to individual participants (**Figure 4C**). Analyses in individual participants identified 5–10 clusters per participant (mean = 7), where each cluster was defined as including more than one sulcus and the cluster count was the maximum number identified among the clustering thresholds. The pairs of sulci within the same group-level cluster were assigned together at multiple thresholds in 80% of participants, on average. We observed the highest degree of agreement for the two LPC clusters – one comprising large sulci (yellow in **Figure 4C**: 94%), and the other including both large sulci and pTS ( medium blue: 91%), followed by the LPFC cluster of pTS that included sulci previously implicated in reasoning (dark blue: 87%) and an LPFC-LPC cluster of pTS (purple: 86%). We observed lower agreement for a large LPFC-LPC cluster including both large sulci and pTS (green: 76%) and a small LPC cluster of pTS comprising only bilateral slocs-v, a pTS (cyan: 70%). Notably, even the lowest of these scores (70%) was significantly higher than the 58% co-assignment for pairs that did not belong to the same group-level cluster (t42 = 10.83, two-tailed p < 0.0001; the lowest value was 19%, between lh.mTOS (an LPC pTS) and lh.lfms (an LPFC pTS). Thus, there was variable agreement between individual and group-level cluster assignments.

Here, we highlight five specific findings based on these group- and individual-based connectivity-based clustering analyses. First, most relevant to the current investigation, the LPFC pTS whose morphological features have been previously linked to individual differences in reasoning performance – right-hemisphere pimfs, pmfs-a, and pmfs-i (Voorhies et al., 2021), along with left pimfs (Willbrand et al., 2022, 2023b, 2024) – tended to cluster together (dark blue cluster in **Figure 4B**). Second, several pTS in LPFC clustered with aipsJ, a pTS in LPC (shown in purple); they may be anatomically connected by the middle branch of the superior longitudinal fasciculus, which develops late (Liang et al., 2022) and is theorized to support communication between the dorsal and ventral attention systems (Thiebaut de Schotten et al., 2011; Parlatini et al., 2017; Suo et al., 2021). This association could be clinically relevant, as the aipsJ is used as a corridor in neurosurgery to reach deeper structures while minimizing damage to other structures (Tomaiuolo and Giordano, 2016; Tomaiuolo et al., 2022). Third, inferior portions of the intraparietal sulcus (IPS-PO; in yellow cluster) clustered separately from more dorsal portions of the IPS, which clustered (shown in red) with the inferior frontal sulcus (ifs). Fourth, the superior frontal sulcus (sfs) tended to cluster with a subset of branches of the superior temporal sulcus (STS) and pTS in inferior frontal cortex (shown in green), and fifth, newly identified pTS in LPC (slocs-v) often clustered by themselves (shown in cyan), although these were the least reliable groupings across participants.

#### Associations with large-scale canonical resting-state fMRI networks

Many sulci in these clusters showed correlations with one or more large-scale canonical resting-state networks in the MIDB Precision Brain Atlas (Hermosillo et al., 2024), consistently across subjects and to a greater degree than for corresponding rotated labels (**Figure 5**). Sulci within the red cluster (large sulci and pTS in LPFC and LPC) were associated with three networks: frontoparietal, dorsal attention, and cingulo-opercular. Sulci in the yellow cluster (large sulci in LPC) correlated most strongly with the dorsal attention and visual networks, and those in the green cluster (large sulci and pTS in LPFC and LPC) correlated most with the default mode network. By contrast, sulci in the dark blue cluster implicated in reasoning (pTS in LPFC), along with the cyan cluster (pTS in LPC) and medium blue cluster (large sulci and pTS in LPFC and LPC), were not clearly associated with any of these large-scale networks, and in the purple cluster (pTS in LPFC and LPC) only two sulci systematically associated with the frontoparietal network. Thus, we see overlap between some, but not all, networks defined based on individual anatomy and large-scale canonical networks. That said, future investigations should explore the correspondence of these sulcal networks and other large-scale networks in the literature, as network definitions and nomenclature vary across the literature (Uddin et al., 2023).

**Figure 5.**
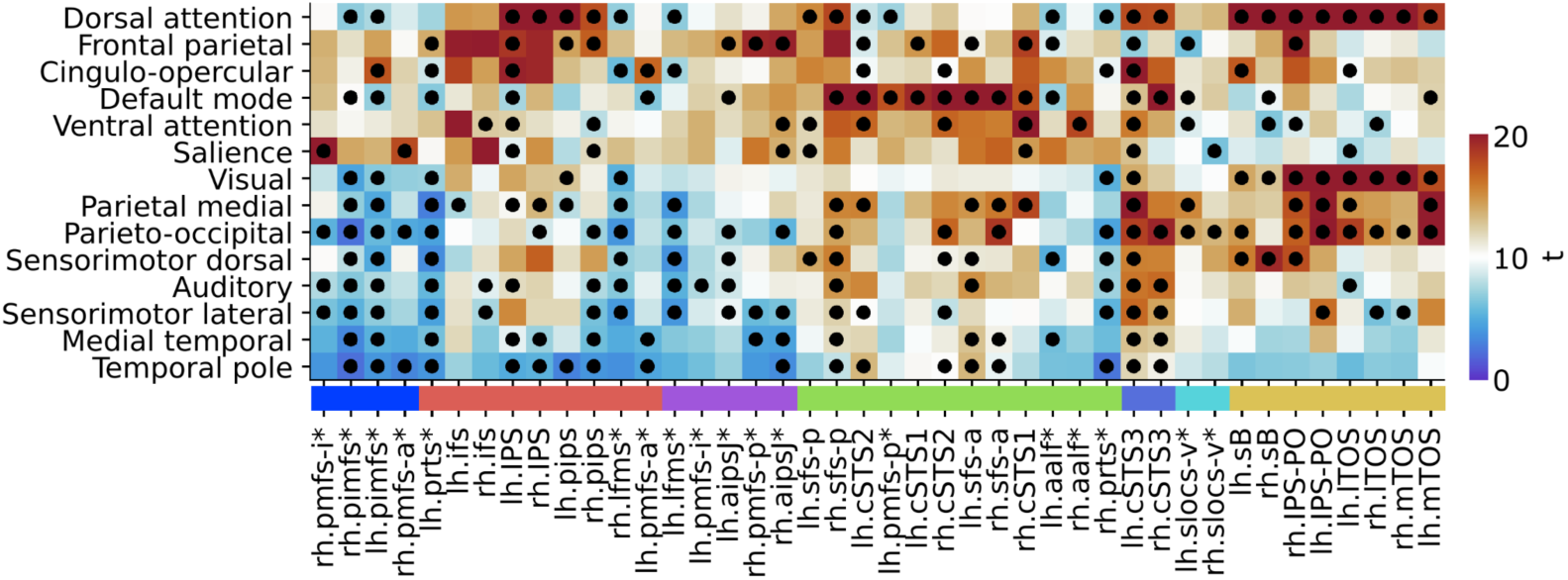
Associations of LPFC and LPC sulci with canonical large-scale resting-state fMRI networks. Correlation strength (compared to zero via a one-sample t test) between each sulcus and each large-scale canonical network (arranged by clusters as in Figure 4B). *Putative tertiary sulcus (pTS). The color scale reflects t-scores ranging from zero to positive values. Dots indicate associations that were significantly more positive/negative for sulci compared to rotated labels (permuted two-tailed p < .05). In general, larger sulci, such as those in the red, green, and yellow clusters, showed pronounced correlations with one or more large-scale networks; by contrast, many pTS, including those in the blue and purple clusters, showed less robust associations with these networks.

#### Addressing a possible confound of spatial autocorrelations

One potential confound is that neighboring sulci might have inflated connectivity due to spatial autocorrelations, known to arise in BOLD fMRI from various physical, physiological, and biological sources that are not all related to functional organization (Burt et al., 2020; Power et al., 2012; Shinn et al., 2023). As such, we controlled for spatial autocorrelations in our main analyses by estimating and regressing out the short-distance positive correlation bias on an individual basis. We found that the clustering patterns could not be explained by spatial autocorrelations among neighboring sulci, for two main reasons. First, the preferential correlation among neighboring sulci in the LPFC pTS cluster (dark blue in **Figure 4B**) was observed despite being penalized via correction for spatial autocorrelations (see **Extended Data** Figure 6-1 for all connectivity fingerprints with and without controlling for spatial autocorrelations). Second, a number of geographically distant sulci clustered together – for example, the pmfs-p and the aipsJ (purple in **Figure 4B**) – demonstrating a novel functional connection between pTS across LPFC and LPC.

The main findings were replicated also when not controlling for spatial autocorrelations (data available upon request). For example, classification accuracies were similarly very high (mean: 96%, range: 70–100%) – albeit not quite as high (t41 = 3.81, p < 0.001). The boost in accuracy with correction for autocorrelations was observed for the sulci that were closest together (< 5 mm geodesic distance) (t971 = 2.43, p < 0.05), as expected, and also for pairs of homologous sulci across the two hemispheres (t20 = 2.98, p < 0.01).

Clustering without control for spatial autocorrelations replicated key findings, e.g. the blue and purple pTS clusters with high co-clustering across participants, but also suggested possible additional distinctions.

### LPFC-LPC sulcal network topology and measures of centrality

We next looked more closely at the connectivity patterns of individual sulci. Perhaps unsurprisingly, sulci had on average the strongest connectivity with other sulci that co-clustered with them (**Figure 6A**, **B**; **Extended Data** Figure 6-1). To identify sulci with relatively high or low network connectivity, we compared measures of centrality for a given node to the average across nodes for that participant. We examined three measures of centrality: degree, betweenness, and participation coefficient (see Materials and Methods for descriptions).

**Figure 6.**
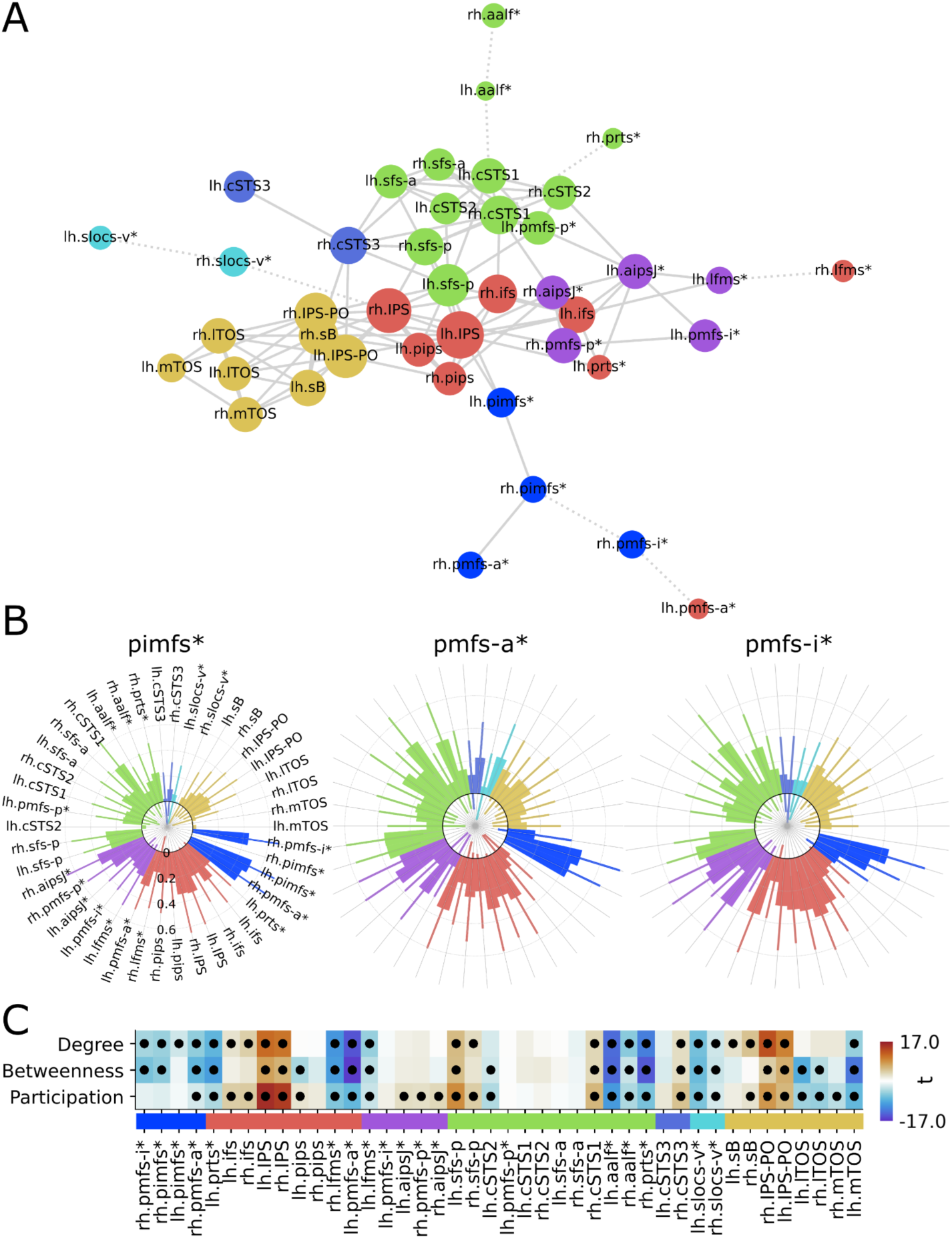
Functional organization of the sulcal LPFC–LPC network. (A) Spring plot of the network derived across participants, showing connections at 0.12 density and in dotted lines the strongest connection for the nodes isolated at this threshold (rh.pmfs-i, lh.pmfs-a, rh.prts, rh.lfms, and bilateral aalf and slocs-v). Each of the five larger clusters shows high connectivity within the cluster. Node size is proportional to mean AUC degree. *Putative tertiary sulcus (pTS). The cluster colors match Figure 4B. (B) Functional connectivity fingerprints (mean, SD) of the right-hemisphere pTS in LPFC under investigation: pimfs, pmfs-i, and pmfs-a (dark blue cluster). For functional connectivity fingerprints of all sulci, with and without correction for spatial autocorrelation, see **Extended Data** Figure 6-1. (C) The nodes associated with significantly higher or lower centrality measures (degree, betweenness, and/or participation coefficient) relative to the mean of values of all nodes for that participant. Cells are shaded according to strength of t-values for each comparison, with warm and cool colors denoting above and below mean. Black dots depict cells with significant (FDR-corrected p < 0.05) results.

Notably, almost all LPFC pTS scored below average with regards to at least one (and often more) centrality measure: bilateral pimfs, pmfs-a, and lfms, aalf, and prts, along with right pmfs-i (**Figure 6C**). By contrast, left pmfs-i showed average centrality values, and right pmfs-p (along with bilateral aipsJ, a pTS in LPC that clustered with it) scored above average on participation coefficient, with trend-level effects for the other centrality measures. These results were very similar when not controlling for spatial autocorrelations (data available upon request). In sum, these results suggest that most LPFC pTS have relatively sparse connectivity; these results can be contrasted with LPC sulci IPS and IPS-PO, which scored consistently high in the centrality metrics across participants (each sulcus p < 0.05, FDR corrected). None of the metrics were linearly associated with head motion (an association was detected for only 1/126 tests: betweenness of rh.ifs was negatively associated with mean FD at β = -2.269, p < 0.03, uncorrected for multiple comparisons). It should be noted that the sulcal network covered large swaths of association cortex relevant for reasoning; these results may not generalize to broader or more granular networks. These graph metrics characterize the network at the group level; individual variability in centrality is explored below in relation to sulcal depth.

Observing that the graph analyses attributed higher centrality to large sulci, such as those in IPS and IPS-PO, than to small ones, such as the pTS in the blue LPFC cluster, we explicitly tested whether functional connectedness was systematically related to sulcal surface area. Specifically, we tested the relation between graph metrics and surface area with linear mixed effect models with random intercepts and slopes for participant and surface area, respectively. We did in fact observe a significant positive association with surface area for all three centrality measures (degree: β = 0.59, p < 0.001; betweenness: β = 0.43, p < 0.001; participation coefficient: β = 0.58, p < 0.001). This relationship could reflect a true feature of brain architecture. Alternatively, it could reflect a methodological confound: given that the fMRI timeseries was averaged over all vertices in a sulcus, it is possible that sulci with a larger surface area had a higher signal-to-noise ratio than smaller ones, perhaps artificially boosting their centrality metrics. Either way, it is all the more noteworthy that many of the small sulci had preferential connectivity to other small sulci rather than the larger ones.

### Sulcal depth and functional connectivity are correlated for specific pTS in LPFC

We tested whether the functional connectivity patterns of right and/or left pimfs, pmfs-a, and pmfs-i sulci measured during performance of a reasoning task would be related to their depth, as we have previously found that the depths of these sulci in the right hemisphere was related to reasoning performance (Voorhies et al., 2021). We found positive associations between depth and centrality measures for left pmfs-i, right pmfs-a, and left pimfs that survived correction for multiple comparisons (**Figure 7A**). These significant results were also obtained when not controlling for spatial autocorrelations (data available upon request).

**Figure 7.**
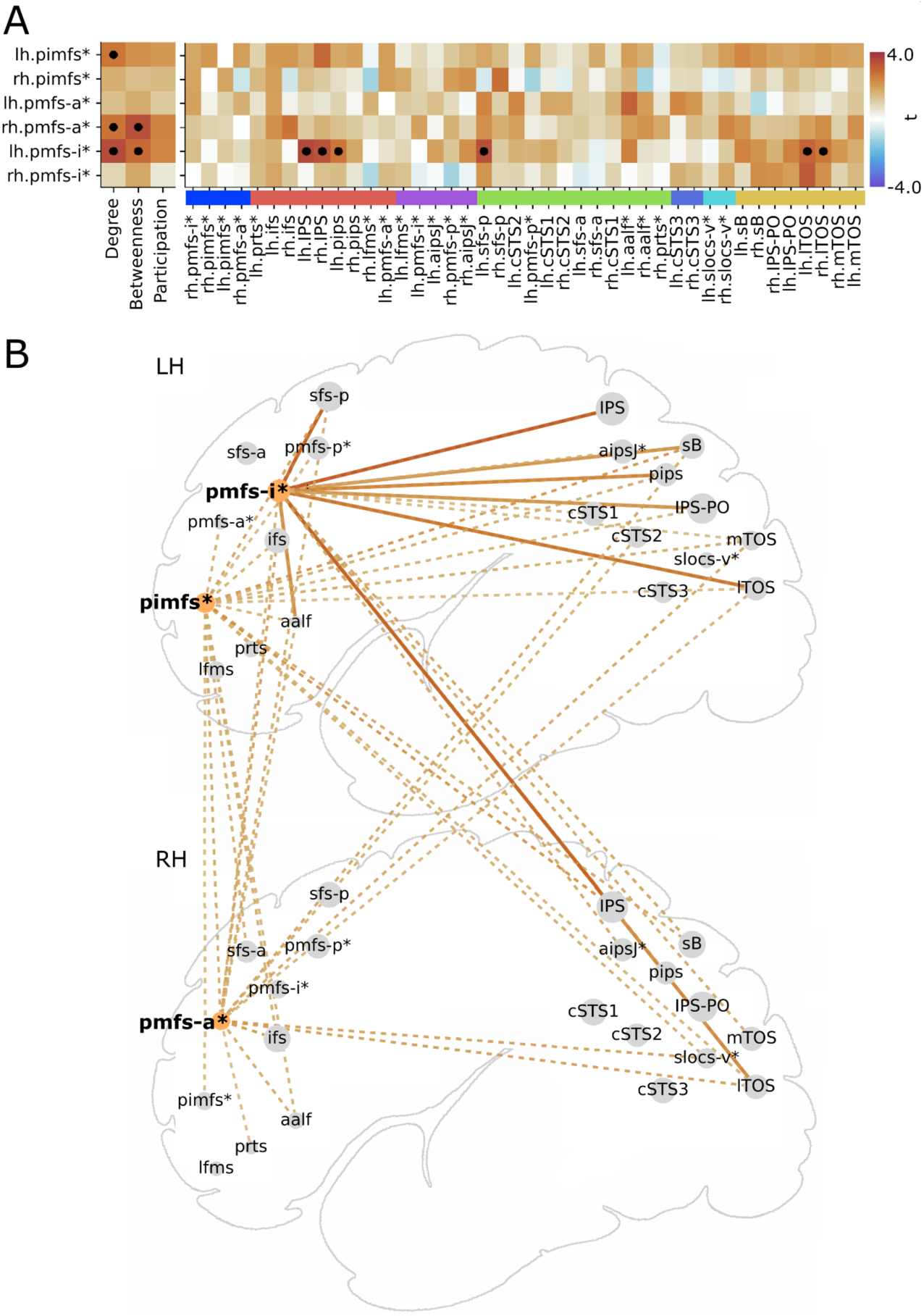
Sulcal depth and functional connectivity are correlated for specific pTS in LPFC. The depths of lh.pmfs-i, rh.pmfs-a, and left pimfs were positively correlated with their functional connectivity, controlling for age in addition to spatial autocorrelations. (A) First three columns: results of regression analyses revealing significant associations between depth and centrality measures for lh.pmfs-i and rh.pmfs-a. Cells are shaded according to strength of t-values for each regression, with warm and cool colors denoting positive and negative correlations, and darker shades indicating higher values. Dots depict cells with significant results (FDR-corrected p < 0.05). Remaining columns: correlations between the depth of each of these sulci and the strength of their pairwise functional connectivity with the other sulci. *Putative tertiary sulcus (pTS). (B) Schematic illustrating the approximate anatomical distribution of sulci showing depth-related increases in pairwise connection strength (solid FDR-corrected, dashed trends at uncorrected p < 0.05). Node size reflects AUC estimate for degree.

**Figure 8.**
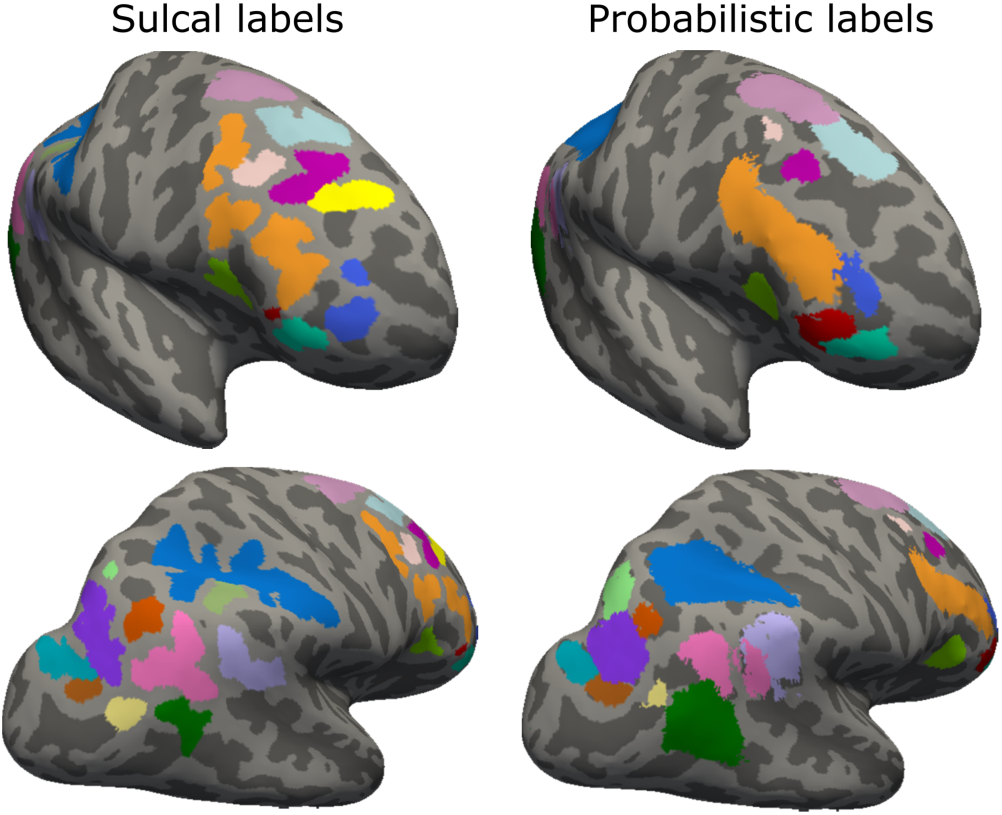
Manual and probabilistic sulcal labels in LPFC (upper row) and LPC (lower row) in an example participant. Group-level probabilistic definitions (minimum 33% overlap in other participants) in LPFC and LPC (except for some pTS such as lh.slocs-v, rh.pmfs-a, and bilateral aipsJ). Label sizes and geodesic distances were also greatly altered between probabilistic and manual definitions. Sulcal colors match Figure 2A.

These depth-centrality relations were not driven by age. There were no significant linear or logarithmic associations between age and either centrality or pairwise connectivity strength (all |β| < 2.26, all FDR-corrected p > 0.36). Likewise, the depth of these sulci was not systematically associated with age (all |β| < 2.65, all positive and negative associations FDR-corrected p > 0.06; see also Willbrand et al., 2023b). Nevertheless, we included a linear effect of age as a covariate in these analyses given the large age range.

We next investigated the spatial pattern of pairwise sulcal connections contributing to the positive depth-connectivity associations shown by graph measures. The edgewise regression analysis revealed that these effects were not driven by connectivity within the LPFC (dark blue) or LPFC-LPC (purple) pTS clusters (**Figure 7B**). Rather, the specific pairwise connections implicated in these depth-connectivity associations included large and deep sulci in LPFC and LPC with high centrality measures (IPS, ifs, sfs-p) and sulci associated with the visual system (IPS-PO, lTOS), but not connectivity to neighboring sulci. Thus left pmfs-i (and, to a lesser extent, right pmfs-a) was more strongly connected across the LPFC–LPC sulcal network in participants for whom these sulci were deeper.

To test the specificity of these results to our target pTS in LPFC, we conducted exploratory analyses testing for sulcal depth-connectivity relationships for three additional pTS of interest: 1) pmfs-p (in green/purple clusters in the left/right hemispheres), an LPFC sulcus that did not cluster with other pmfs components and has not – unlike them – been linked to reasoning performance, 2) aipsJ (in the LPFC-LPC purple cluster), a well-known pTS in LPC (Zlatkina and Petrides, 2014), and 3) slocs-v (small cyan cluster), a newly identified pTS in LPC that is variably present and has been associated with visuospatial perception (Willbrand et al., 2023c). These exploratory analyses revealed several significant depth–connectivity associations, with deeper sulci associated with stronger connectivity (bilateral aipsJ: degree and participation coefficient (β > 2.71, FDR-corrected p < 0.05; rh.slocs-v: all three metrics (β > 3.14, FDR-corrected p > 0.05; figure available upon request). Thus, positive associations between sulcal depth and network centrality were not unique to target pTS in LPFC, but could also be found in several pTS in LPC.

### Variable correspondence between manually defined and probabilistic sulcal labels

We created probabilistic labels based on spatial overlap across at least 33% of participants. Despite this liberal threshold, four sulci – all pTS – did not meet this criterion (lh.slocs-v, rh.pmfs-a, and bilateral aipsJ). For the 38/42 sulci with a probabilistic definition, strength of pairwise functional connectivity was moderately correlated with manual labels at the individual level (Pearson r = 0.58, pooling all participants and connections). The correlation between sulcal fingerprints was on average 0.70, but ranged from -0.51 to 0.98 across sulci and participants. In other words, the results for probabilistic labels were in some cases negatively correlated with manual ones; thus, they did not represent the same functional regions and network connectivity as the manual definitions in individual participants.

## Discussion

This study presents a novel approach of defining sulcal functional connectomes, building on prior work involving seeded functional connectivity based on specific sulci (Amiez et al., 2023; Ducret et al., 2024; Lopez-Persem et al., 2019) and associations between individual sulci and canonical rs-fMRI networks (Miller et al., 2021; Willbrand et al., 2023a, 2023c). Here, we characterize individual LPFC-LPC sulcal connectomes, including smaller sulci that are typically overlooked (referred to here as pTS) in the context of a pediatric study of reasoning. We show that sulci have differentiable connectivity fingerprints, but could nevertheless be grouped together based on similar patterns of connectivity. Further, we demonstrate, via control analyses involving rotated sulcal labels and group-derived probabilistic labels, that network connectivity is meaningfully related to individual sulcal anatomy. We also report, for the first time (to our knowledge), associations between individual variability in sulcal depth and functional connectivity between LPFC and LPC – consistent with the hypothesis posited decades ago that sulcal morphology, including that of pTS in association cortices, is relevant to brain function (Sanides, 1962, 1964).

### Sulcal classification and clustering based on functional connectivity profiles

Classification analyses revealed that most sulci had characteristic functional connectivity profiles, with mean pairwise classification accuracy of 96%. Based on similarity of their connectivity patterns, however, the sulci formed clusters of various configurations. Using the group-derived clusters as a point of departure for exploring individual-level clustering, we observed above-chance, but variable, agreement between group-level and individual-level cluster assignments, speaking to individual variability that is important to characterize.

Of note, the two clusters (dark blue and purple in **Figure 4B**) consisting only of pTS did not map clearly onto any of the large-scale canonical networks considered here (Hermosillo et al., 2024). The dark blue cluster included sulci previously linked to reasoning performance (Voorhies et al., 2021; Willbrand et al., 2023a, 2023b). We observed strong local connectivity among these sulci, and particularly weak connectivity between them and LPC sulci, consistent with local recurrent connectivity in support of higher-level cognition. This sulcal connectivity may be mediated by short-range fiber tracts in superficial white matter (Van Essen et al., 2013; Reveley et al., 2015) whose careful characterization and quantification awaits (Schilling et al., 2023).

The sulcal-functional dissociations identified here are complementary to the handful of recent studies that have compared connectivity patterns of manually defined sulci in LPFC and LPC. One study showed that the neighboring pTS pmfs-a and pmfs-p overlapped with different large-scale networks (Miller et al., 2021). Another study, in macaques, showed different clustering based on whole-brain functional connectivity for the putative homologs of pmfs-a and pmfs-i (Amiez et al., 2023). Integrating these parallel tracks of research, and noting that pimfs, pmfs-a, pmfs-i, and pmfs-p are roughly positioned along a rostral-caudal axis, differences in their functional connectivity patterns may reflect their geographic location within a larger anatomical and functional hierarchical gradient in LPFC (Miller et al., 2021). Further study is required to test the contributions of morphology vs. topographic location on the functional connectivity patterns of these and other sulci.

### Sulci as a coordinate system for functional connectivity analyses

A robust literature shows that cortical areas and large-scale networks can be defined in vivo based on patterns of functional connectivity (Gordon et al., 2017; Hermosillo et al., 2024; Power et al., 2011; Yeo et al., 2011). However, areal and network definitions vary markedly among individuals (Seitzman et al., 2019; Gordon and Nelson, 2021; Smith et al., 2021), and may differ, at least in subtle ways, across the lifespan (e.g., Cui et al., 2020; Tooley et al., 2022) and clinical populations (e.g., Lynch et al., 2024; Persichetti et al., 2024) – and as a function of analytic approach (Braga and Buckner, 2017; Cookson and D’Esposito, 2023; Dixon et al., 2018; Gordon et al., 2020; Kwon et al., 2025; Luckett et al., 2023; Uddin et al., 2023).

To address this variability, we propose to use sulci as a personalized coordinate space, building on prior work and theorizing (Ducret et al., 2024; Miller and Weiner, 2022; Lopez-Persem et al., 2019; Sun et al., 2016). While a previous study found no correspondence between functional parcellations and major sulci defined in atlases (Zhi et al., 2022), we encourage further investigation at the individual level – including the numerous smaller, morphologically variable pTS, many of which do not appear in common atlases. Having observed inconsistencies in functional connectomes between group-derived probabilistic sulcal labels and individual ones, we propose that probabilistic definitions should be employed, or at least interpreted, with caution. Moreover, probabilistic labels cannot *yet* be used to derive accurate morphological sulcal metrics or test sulcal associations with white matter or behavior. That said, the probabilistic labels derived from this dataset could (i) serve as a starting point for trainees undertaking manual labeling of these sulci, and (ii) be used to characterize the location and inter-individual variability in the definition of these sulci (GitHub link to be provided). Ongoing work using deep-learning approaches suggests that probabilistic definitions could be used to derive accurate morphological sulcal metrics in the near future (Borne et al., 2020; Lee et al., 2024; Lyu et al., 2021). Moreover, identifying sulci requires only a short anatomical (T1) scan, as compared with many runs of task-based, resting-state, or multimodal data aimed at identifying cortical areas for network analyses.

### Network centrality is correlated with sulcal depth

Relative to the full set of sulci in the LPFC-LPC network, none of the pTS showed high centrality. From a network perspective, this is consistent with the “preferential age attachment” theory, which suggests that early-developing nodes should become stronger hubs than those born later (Barabási and Albert, 1999; Diez et al., 2022). Notably, although centrality was low on average for pTS, it was—for half of the pTS tested, both in our main analyses on LPFC sulci and exploratory ones on LPC sulci—higher for individuals with deeper pTS, driven by stronger and more widespread connectivity across the LPFC-LPC network. Given that sulcal development during gestation has been theorized to serve as the foundation of functional architecture and cognition (Régis et al., 2005; Cachia et al., 2021; Weiner, 2023), the hypothesis that the overall low, but variable, centrality of pTS relates to later neural and cognitive development should be tested longitudinally across early development.

Further, while we showed that the depth of the left pimfs was correlated with a measure of network centrality, we combined the dorsal and ventral branches of this sulcus to maximize our sample size, as in our previous study relating sulcal depth to reasoning (Voorhies et al., 2021). However, having shown that the presence of the ventral branch of the left pimfs was associated with reasoning performance in both pediatric and adult samples (Willbrand et al., 2022, 2024), exploring how functional connectivity differs in the presence or absence of left pimfs-v would serve as an effective testbed for continuing to explore how functional connectivity relates to sulcal presence (Amiez and Petrides, 2014; Artiges et al., 2006; Lopez-Persem et al., 2019; Wilbrand et al., 2023a).

## Limitations and future directions

This study should be considered an early step in a new line of inquiry, with limitations to be addressed in further research. First, the amount of fMRI data per participant, while fairly common in developmental samples, is lower than recommended to achieve high test-retest reliability for individual connections (Birn et al., 2013; Elliott et al., 2019; Laumann et al., 2015; Noble et al., 2017). Second, as this study builds on prior findings in a project examining the neural basis of reasoning during development, it focuses on a specific dataset. Thus, it is an open question as to whether the results would generalize to other sulci, or other datasets involving other age groups or types of fMRI data – including resting-state fMRI data (but see Cole et al., 2016; Gratton et al., 2018; Salvo et al., 2021). Third, there may be residual head motion effects on connectivity. Fourth, the small sample size (or the inclusion of head motion as a covariate at the group level) likely explains the lack of effects of age on functional connectivity that have been widely documented (Grayson and Fair, 2017; Luo et al., 2024; Uddin et al., 2011), and perhaps also on sulcal depth. However, this does not detract from our individual differences analysis showing that depth and connectivity were correlated with one another – albeit not age – for a subset of sulci across a heterogeneous sample.

Limitations aside, we have made a number of observations in this foundational study that lead us to make several predictions for studies involving other datasets: 1) sulci can be discriminated based on their connectivity patterns, outperforming null models, 2) some sulci would show similar patterns of connectivity despite their size and geographic distance, and 3) centrality would, for some sulci, vary across individuals as a function of depth. The present results also prompt further multi-modal characterization of pTS – including white matter measures, which may in some cases serve as a mechanistic link between cortical folding and functional brain architecture (Bouhali et al., 2024; Kruggel and Solodkin, 2023).

## Conflict of interest statement

The authors declare no competing financial interests.

### Acknowledgments

This research was supported by NICHD R21HD100858 (Weiner, Bunge), NSF CAREER Award 2042251 (Weiner), and NIMH R01MH133637 (Bunge). Funding for the original data collection and curation was provided by NINDS R01 NS057156 (Bunge, Ferrer) and NSF BCS1558585 (Bunge, Wendelken). EHW was supported by the Medical Scientist Training Program Grant T32 GM140935 (Willbrand). We thank Allison Chen for assistance with manuscript preparation.

## Extended Data

**Extended Data Figure 2-1.**
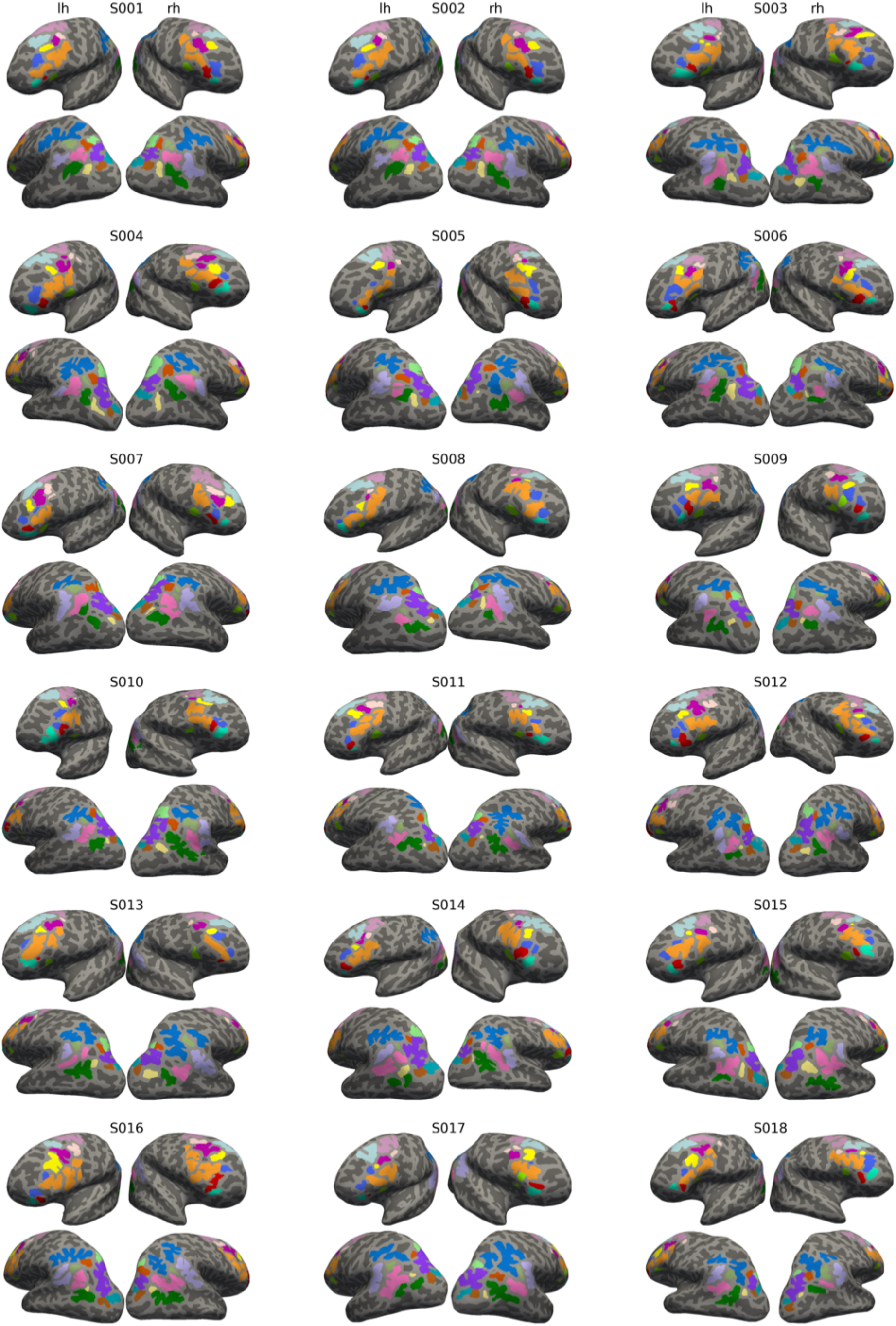
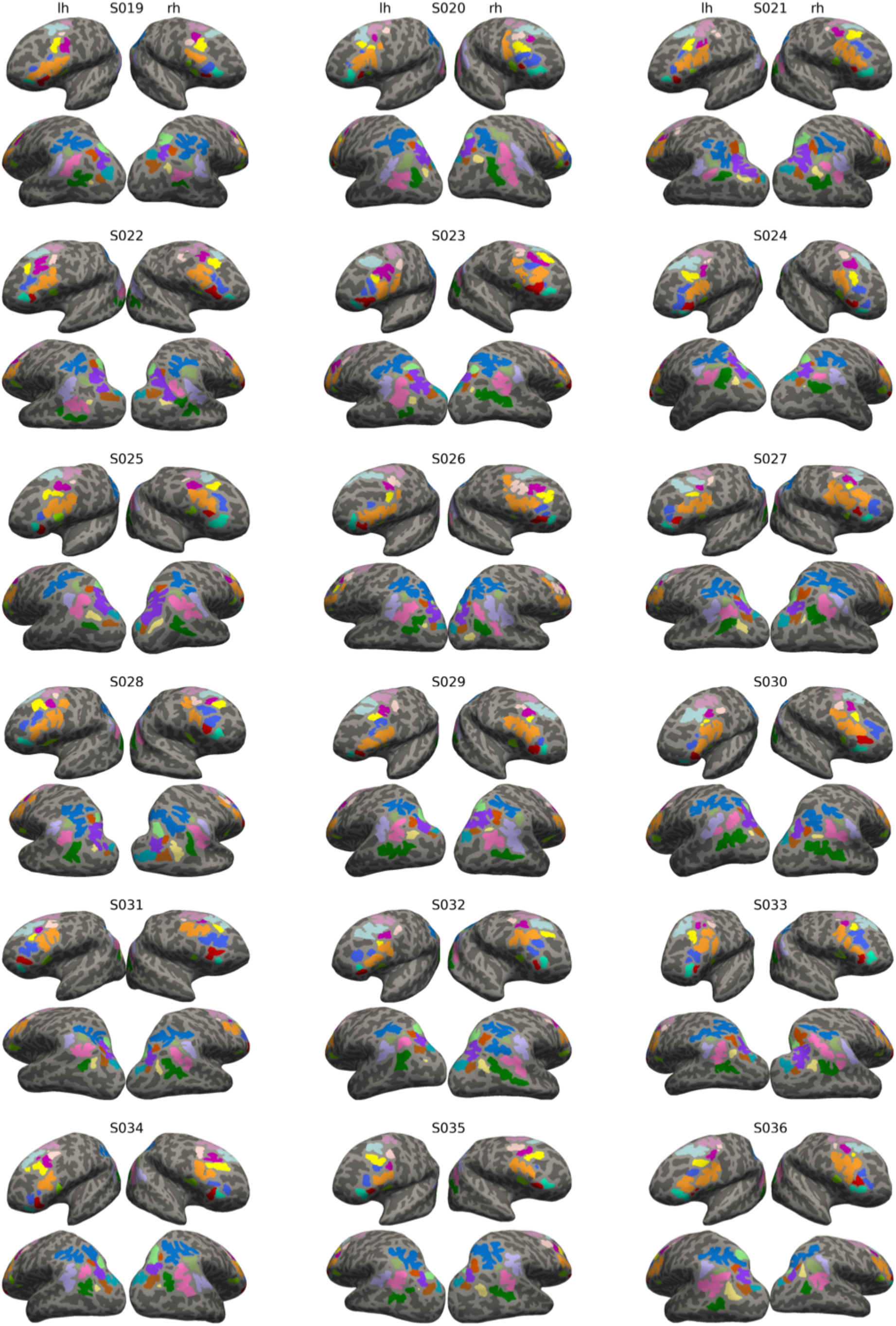
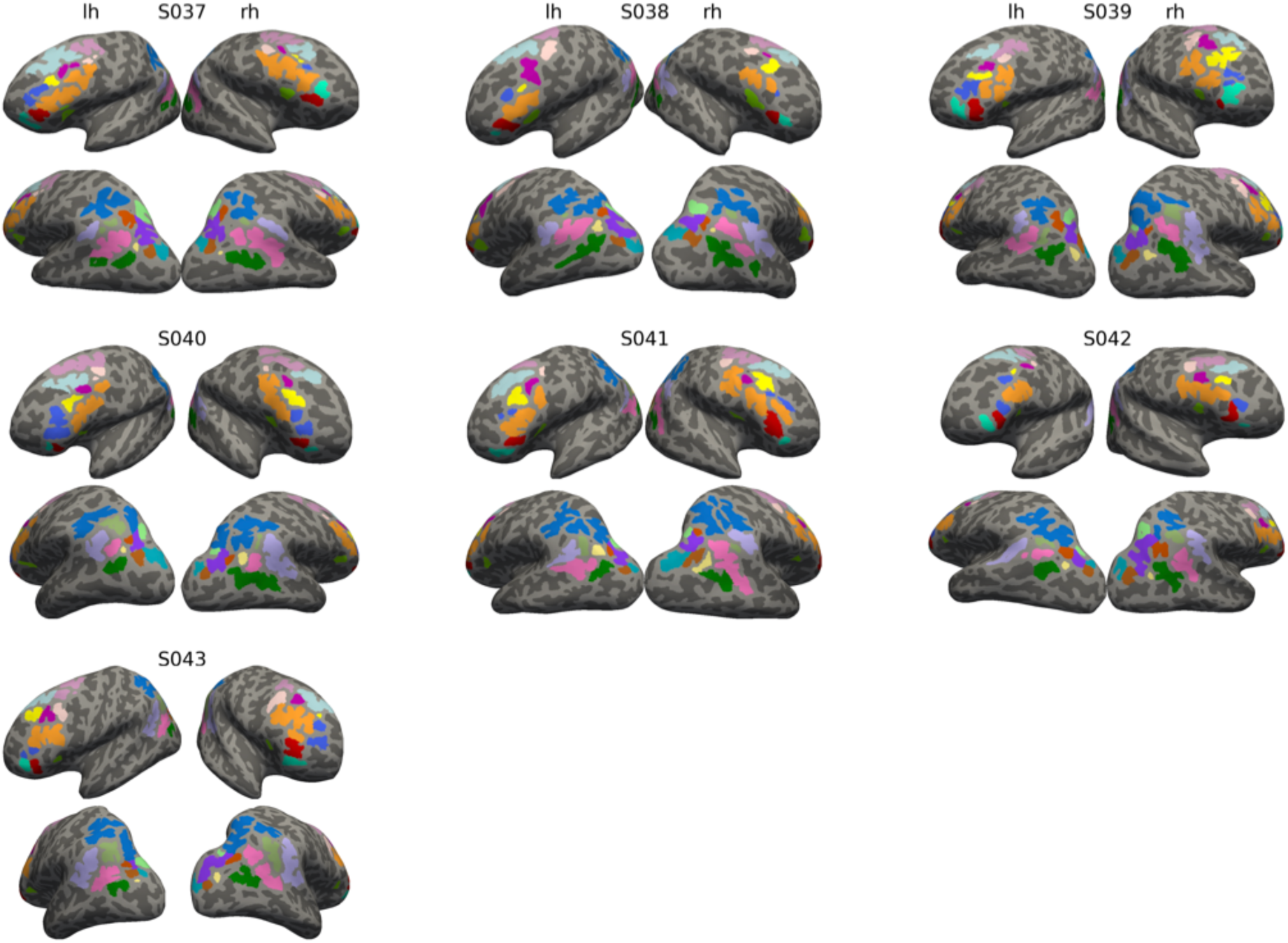
Sulcal definitions in all participants. Colors and sulcal names match Figure 2A.

**Extended Data Figure 6-1.**
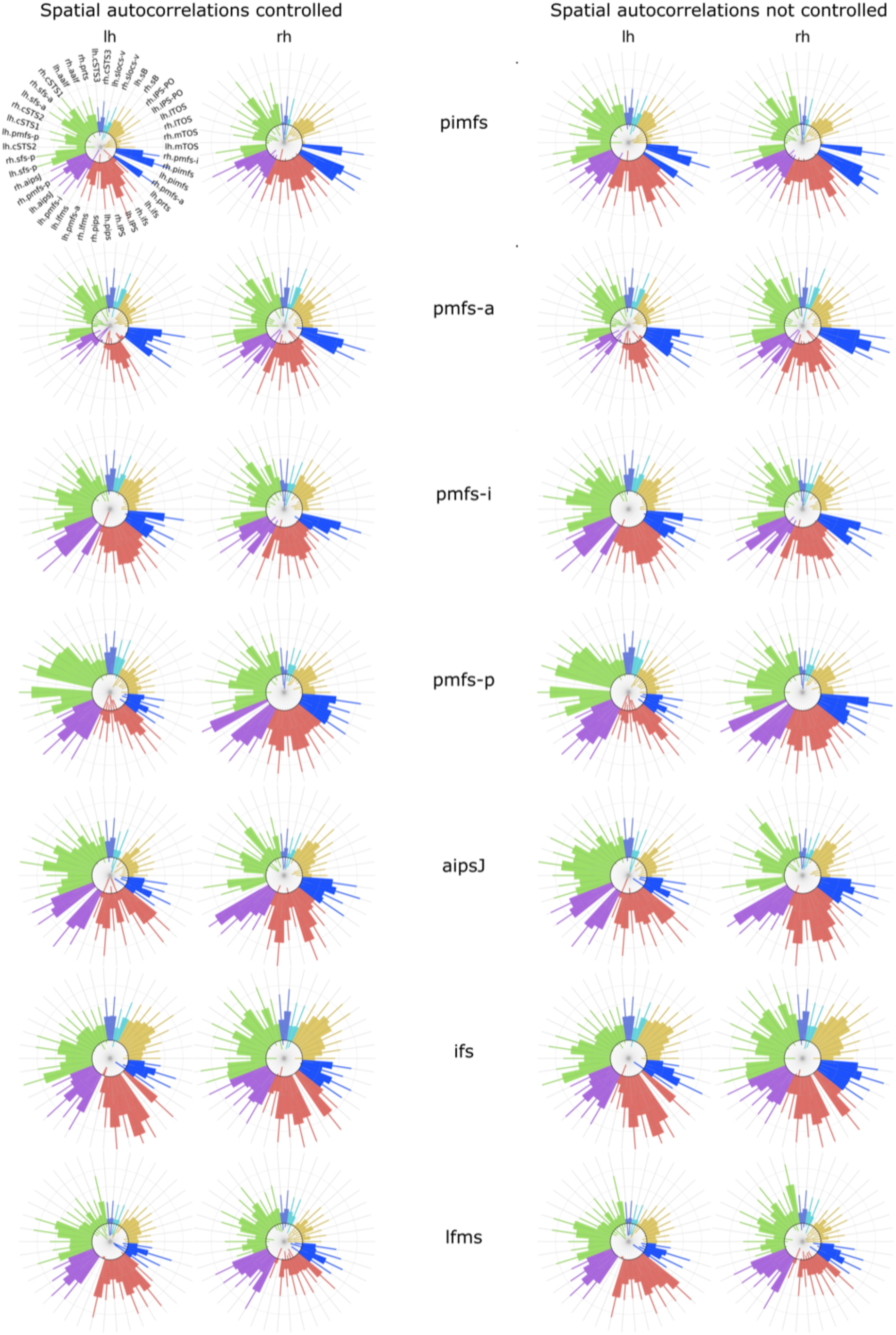
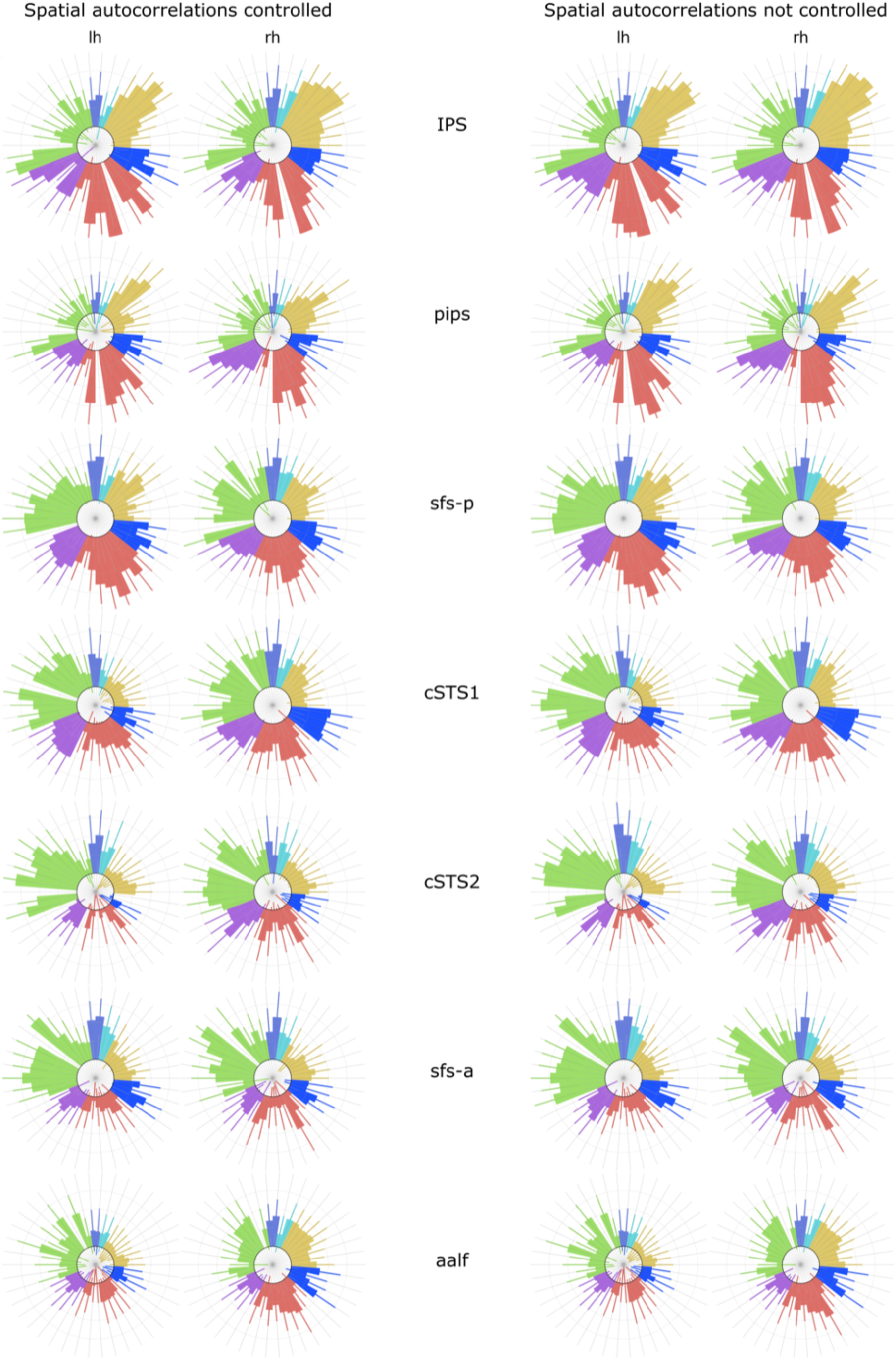
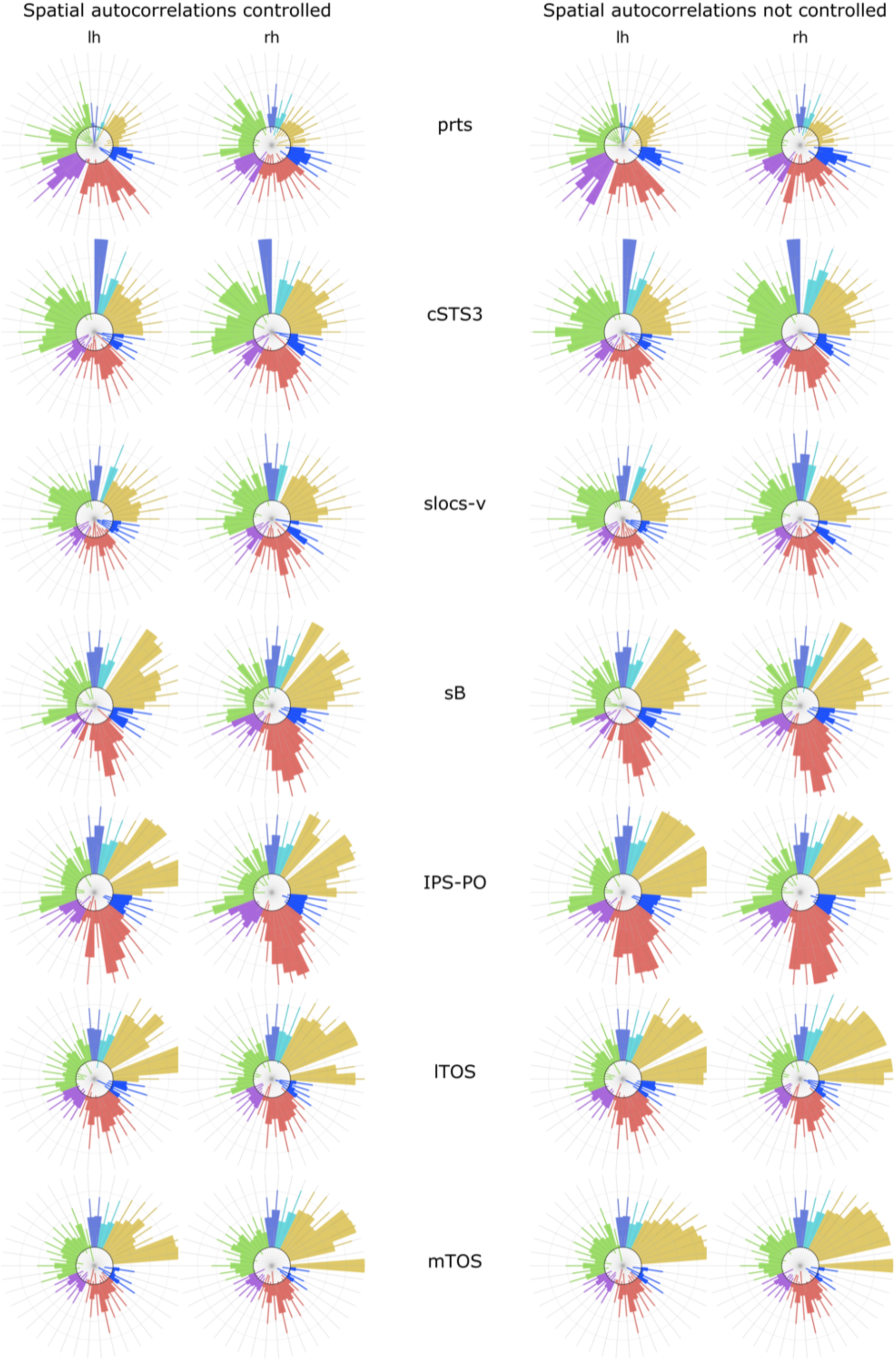
Functional connectivity fingerprints with and without correction for spatial autocorrelations. The colors match the clusters in Figure 4B.

## References

1. Aichelburg C, Urbanski M, Thiebaut de Schotten M, Humbert F, Levy R, Volle E (2016) Morphometry of left frontal and temporal poles predicts analogical reasoning abilities. Cereb Cortex 26:915–932.

2. Alexander-Bloch AF, Shou H, Liu S, Satterthwaite TD, Glahn DC, Shinohara RT, Vandekar SN, Raznahan A (2018) On testing for spatial correspondence between maps of human brain structure and function. NeuroImage 178: 540–551.

3. Amiez C, Neveu R, Warrot D, Petrides M, Knoblauch K, Procyk E (2013) The location of feedback-related activity in the midcingulate cortex is predicted by local morphology. J Neurosci 33:2217–2228.

4. Amiez C, Petrides M (2014) Neuroimaging evidence of the anatomo-functional organization of the human cingulate motor areas. Cereb Cortex 24:563–578.

5. Amiez C, Sallet J, Giacometti C, Verstraete C, Gandaux C, Morel-Latour V, Meguerditchian A, Hadj-Bouziane F, Ben Hamed S, Hopkins WD, Procyk E, Wilson CRE, Petrides M (2023) A revised perspective on the evolution of the lateral frontal cortex in primates. Sci Adv 9:eadf9445.

6. Armstrong E, Schleicher A, Omran H, Curtis M, Zilles K (1995) The ontogeny of human gyrification. Cereb Cortex 5:56–63.

7. Artiges E, Martelli C, Naccache L, Bartrés-Faz D, LeProvost JB, Viard A, Paillère-Martinot ML, Dehaene S, Martinot JL (2006) Paracingulate sulcus morphology and fMRI activation detection in schizophrenia patients. Schizophr Res 82:143–151.

8. Barabási AL, Albert R (1999) Emergence of scaling in random networks. Science 286:509– 512.

9. Behzadi Y, Restom K, Liau J, Liu TT (2007) A component based noise correction method (CompCor) for BOLD and perfusion based fMRI. NeuroImage 37:90–101.

10. Benjamini Y, Hochberg Y (1995) Controlling the false discovery rate: a practical and powerful approach to multiple testing. J R Stat Soc Ser B Methodol 57:289–300.

11. Birn RM, Molloy EK, Patriat R, Parker T, Meier TB, Kirk GR, Nair VA, Meyerand ME, Prabhakaran V (2013) The effect of scan length on the reliability of resting-state fMRI connectivity estimates. NeuroImage 83:550–558.

12. Bodin C, Takerkart S, Belin P, Coulon O (2018) Anatomo-functional correspondence in the superior temporal sulcus. Brain Struct Funct 223:221–32.

13. Borne L, Rivière D, Mancip M, Mangin JF (2020) Automatic labeling of cortical sulci using patch- or CNN-based segmentation techniques combined with bottom-up geometric constraints. Med Image Anal 62:101651.

14. Borst G, Cachia A, Vidal J, Simon G, Fischer C, Pineau A, Poirel N, Mangin JF, Houdé O (2014) Folding of the anterior cingulate cortex partially explains inhibitory control during childhood: a longitudinal study. Dev Cogn Neurosci 9:126–35.

15. Bouhali F, Dubois J, Hoeft F, Weiner KS (2024) Unique longitudinal contributions of sulcal interruptions to reading acquisition in children. eLife 13:RP103007.

16. Braga RM, Buckner RL (2017) Parallel interdigitated distributed networks within the individual estimated by intrinsic functional connectivity. Neuron 95:457–471.e5.

17. Brett M et al. (2022) nipy/nibabel: (4.0.0). doi:10.5281/zenodo.6658382

18. Burt JB, Helmer M, Shinn M, Anticevic A, Murray JD (2020) Generative modeling of brain maps with spatial autocorrelation. NeuroImage 220:117038.

19. Cachia A, Borst G, Jardri R, Raznahan A, Murray GK, Mangin JF, Plaze M (2021) Towards deciphering the fetal foundation of normal cognition and cognitive symptoms from sulcation of the cortex. Front Neuroanat 15:712862.

20. Cachia A, Roell M, Mangin JF, Sun ZY, Jobert A, Braga L, Houde O, Dehaene S, Borst G (2018) How interindividual differences in brain anatomy shape reading accuracy. Brain Struct Funct 223:701–712.

21. Chi JG, Dooling EC, Gilles FH (1977) Gyral development of the human brain. Ann Neurol 1:86–93.

22. Cole MW, Ito T, Bassett DS, Schultz DH (2016) Activity flow over resting-state networks shapes cognitive task activations. Nat Neurosci 19:1718–1726.

23. Cookson SL, D’Esposito M (2023) Evaluating the reliability, validity, and utility of overlapping networks: Implications for network theories of cognition. Hum Brain Mapp 44:1030–1045.

24. Cordeau M, Bichoutar I, Meunier D, Loh KK, Michaud I, Coulon O, Auzias G, Belin P (2023) Anatomo-functional correspondence in the voice-selective regions of human prefrontal cortex. NeuroImage. 279:120336.

25. Cox RW (1996) AFNI: Software for analysis and visualization of functional magnetic resonance neuroimages. Comput Biomed Res 29:162–173.

26. Cui Z, Li H, Xia CH, Larsen B, Adebimpe A, Baum GL, Cieslak M, Gur RE, Gur RC, Moore TM, Oathes DJ, Alexander-Bloch AF, Raznahan A, Roalf DR, Shinohara RT, Wolf DH, Davatzikos C, Bassett DS, Fair DA, Fan Y, Satterthwaite TD (2020) Individual variation in functional topography of association networks in youth. Neuron 106:340– 53.

27. Dale AM, Fischl B, Sereno MI (1999) Cortical surface-based analysis: I. segmentation and surface reconstruction. NeuroImage 9:179–194.

28. Derrfuss J, Brass M, von Cramon DY, Lohmann G, Amunts K (2009) Neural activations at the junction of the inferior frontal sulcus and the inferior precentral sulcus: interindividual variability, reliability, and association with sulcal morphology. Hum Brain Mapp 30:299–311.

29. Destrieux C, Fischl B, Dale A, Halgren E (2010) Automatic parcellation of human cortical gyri and sulci using standard anatomical nomenclature. NeuroImage 53:1–15.

30. Di Carlo DT, Filice ME, Fava A, Quilici F, Fuochi B, Cecchi P, Donatelli G, Restani L, Nardini V, Turillazzi E, Cosottini M, Perrini P (2023) Development of associational fiber tracts in fetal human brain: a cadaveric laboratory investigation. Brain Struct Funct 228:2007–2015.

31. Diez I, Garcia-Moreno F, Carral-Sainz N, Stramaglia S, Nieto-Reyes A, D’Amato M, Cortes JM, Bonifazi P (2022) Linking hubness, embryonic neurogenesis, transcriptomics and diseases in human brain networks. bioRxiv 2022.04.01.486541; doi:10.1101/2022.04.01.486541

32. Dixon ML, De La Vega A, Mills C, Andrews-Hanna J, Spreng RN, Cole MW, Christoff K (2018) Heterogeneity within the frontoparietal control network and its relationship to the default and dorsal attention networks. Proc Natl Acad Sci U S A 115:E1598–E1607.

33. Eichert N, Watkins KE, Mars RB, Petrides M (2021) Morphological and functional variability in central and subcentral motor cortex of the human brain. Brain Struct Funct 226:263–279.

34. Elliott ML, Knodt AR, Cooke M, Kim MJ, Melzer TR, Keenan R, Ireland D, Ramrakha S, Poulton R, Caspi A, Moffitt TE, Hariri AR (2019) General functional connectivity: Shared features of resting-state and task fMRI drive reliable and heritable individual differences in functional brain networks. NeuroImage 189:516–532.

35. Esteban O, Birman D, Schaer M, Koyejo OO, Poldrack RA, Gorgolewski KJ (2017) MRIQC: Advancing the automatic prediction of image quality in MRI from unseen sites. PLOS ONE 12:e0184661.

36. Esteban O, Markiewicz CJ, Blair RW, Moodie CA, Isik AI, Erramuzpe A, Kent JD, Goncalves M, DuPre E, Snyder M, Oya H, Ghosh SS, Wright J, Durnez J, Poldrack RA, Gorgolewski KJ (2019) fMRIPrep: a robust preprocessing pipeline for functional MRI. Nat Methods 16:111–116.

37. Fedeli D, Del Maschio N, Del Mauro G, Defendenti F, Sulpizio S, Abutalebi J (2022) Cingulate cortex morphology impacts on neurofunctional activity and behavioral performance in interference tasks. Sci Rep 12:13684.

38. Fischl B, Sereno MI, Tootell RBH, Dale AM (1999) High-resolution intersubject averaging and a coordinate system for the cortical surface. Hum Brain Mapp 8:272–284.

39. Fischl B (2004) Automatically parcellating the human cerebral cortex. Cereb Cortex 14:11– 22.

40. Fischl B, Dale AM (2000) Measuring the thickness of the human cerebral cortex from magnetic resonance images. Proc Natl Acad Sci 97:11050–11055.

41. Friston KJ, Williams S, Howard R, Frackowiak RSJ, Turner R (1996) Movement-related effects in fMRI time-series: movement artifacts in fMRI. Magn Reson Med 35:346– 355.

42. Ducret M, Giacometti C, Dirheimer M, Dureux A, Autran-Clavagnier D, Hadj-Bouziane F, Verstraete C, Lamberton F, Wilson CR, Amiez C, Procyk E (2024) Medial to lateral frontal functional connectivity mapping reveals the organization of cingulate cortex. Cereb Cortex 34:bhae322.

43. Galar M, Fernández A, Barrenechea E, Bustince H, Herrera F (2011) An overview of ensemble methods for binary classifiers in multi-class problems: Experimental study on one-vs-one and one-vs-all schemes. Pattern Recognit 44:1761–1776.

44. Garrison JR, Fernyhough C, McCarthy-Jones S, Haggard M, Australian Schizophrenia Research Bank, Simons JS (2015) Paracingulate sulcus morphology is associated with hallucinations in the human brain. Nat Commun 6:8956.

45. Germann J, Chakravarty MM, Collins DL, Petrides M (2020) Tight coupling between morphological features of the central sulcus and somatomotor body representations: a combined anatomical and functional MRI study. Cereb Cortex 30:1843–1854.

46. Gordon EM, Laumann TO, Gilmore AW, Newbold DJ, Greene DJ, Berg JJ, Ortega M, Hoyt-Drazen C, Gratton C, Sun H, Hampton JM, Coalson RS, Nguyen AL, McDermott KB, Shimony JS, Snyder AZ, Schlaggar BL, Petersen SE, Nelson SM, Dosenbach NUF (2017) Precision functional mapping of individual human brains. Neuron 95:791–807.

47. Gordon EM, Laumann TO, Marek S, Raut RV, Gratton C, Newbold DJ, Greene DJ, Coalson RS, Snyder AZ, Schlaggar BL, Petersen SE, Dosenbach NUF, Nelson SM (2020) Default-mode network streams for coupling to language and control systems. Proc Natl Acad Sci U S A 117:17308–17319.

48. Gordon EM, Nelson SM (2021) Three types of individual variation in brain networks revealed by single-subject functional connectivity analyses. Curr Opin Behav Sci 40:79–86.

49. Gorgolewski K, Burns CD, Madison C, Clark D, Halchenko YO, Waskom ML, Ghosh SS (2011) Nipype: a flexible, lightweight and extensible neuroimaging data processing framework in Python. Front Neuroinformatics 5.

50. Gratton C, Laumann TO, Nielsen AN, Greene DJ, Gordon EM, Gilmore AW, Nelson SM, Coalson RS, Snyder AZ, Schlaggar BL, Dosenbach NUF, Petersen SE (2018) Functional brain networks are dominated by stable group and individual factors, not cognitive or daily variation. Neuron 98:439–452.e5.

51. Grayson DS, Fair DA (2017) Development of large-scale functional networks from birth to adulthood: A guide to the neuroimaging literature. NeuroImage 160:15–31.

52. Greve DN, Fischl B (2009) Accurate and robust brain image alignment using boundary-based registration. NeuroImage 48:63–72.

53. Hagberg AA, Schult DA, Swart PJ (2008) Exploring network structure, dynamics, and function using NetworkX. In *Proceedings of the 7th Python in Science Conference* (eds Varoquaux G, Vaught T, Millman J), 11–15.

54. Harris CR et al. (2020) Array programming with NumPy. Nature 585:357–362.

55. Hermosillo RJM, Moore LA, Feczko E, Miranda-Domínguez O, Pines A, Dworetsky A, Conan G, Mooney MA, Randolph A, Graham A, Adeyemo B, Earl E, Perrone A, Carrasco CM, Uriarte-Lopez J, Snider K, Doyle O, Cordova M, Koirala S, Grimsrud GJ, Byington N, Nelson SM, Gratton C, Petersen S, Feldstein Ewing SW, Nagel BJ, Dosenbach NUF, Satterthwaite TD, Fair DA (2024) A precision functional atlas of personalized network topography and probabilities. Nat Neurosci 27:1000–1013.

56. Hunter JD (2007) Matplotlib: A 2D Graphics Environment. Comput Sci Eng 9:90–95.

57. Huster RJ, Enriquez-Geppert S, Pantev C, Bruchmann M (2014) Variations in midcingulate morphology are related to ERP indices of cognitive control. Brain Struct Funct 219, 49–60.

58. Hwang K, Bertolero MA, Liu WB, D’Esposito M (2017) The human thalamus is an integrative hub for functional brain networks. J Neurosci 37: 5594–5607.

59. Im K, Grant PE (2019) Sulcal pits and patterns in developing human brains. NeuroImage 185:881–890.

60. Jenkinson M, Bannister P, Brady M, Smith S (2002) Improved optimization for the robust and accurate linear registration and motion correction of brain images. NeuroImage 17:825–841.

61. Kamada T, Kawai S (1989) An algorithm for drawing general undirected graphs. Inf Process Lett 31.1:7–15.

62. Klein A, Ghosh SS, Bao FS, Giard J, Häme Y, Stavsky E, Lee N, Rossa B, Reuter M, Chaibub Neto E, Keshavan A (2017) Mindboggling morphometry of human brains. PLOS Comput Biol 13:e1005350.

63. Krawczyk DC, Michelle McClelland M, Donovan CM (2011) A hierarchy for relational reasoning in the prefrontal cortex. Cortex 47:588–597.

64. Kruggel F, Solodkin A (2023) Gyral and sulcal connectivity in the human cerebral cortex. Cereb Cortex 33:4216–4229.

65. Kwon YH, Salvo JJ, Anderson NL, Edmonds D, Holubecki AM, Lakshman M, Yoo K, Yeo BT, Kay K, Gratton C, Braga RM (2025) Situating the salience and parietal memory networks in the context of multiple parallel distributed networks using precision functional mapping. Cell Rep 44:115207.

66. Laumann TO, Gordon EM, Adeyemo B, Snyder AZ, Joo SJ, Chen MY, Gilmore AW, McDermott KB, Nelson SM, Dosenbach NUF, Schlaggar BL, Mumford JA, Poldrack RA, Petersen SE (2015) Functional system and areal organization of a highly sampled individual human brain. Neuron 87:657–670.

67. Lee S, Lee S, Willbrand EH, Parker BJ, Bunge SA, Weiner KS, Lyu I (2024) Leveraging input-level feature deformation with guided-attention for sulcal labeling. IEEE Trans Med Imaging, doi:10.1109/TMI.2024.3468727.

68. Li S, Shan Y, Qiao H, Chu L, Zeng D, Zhao T, Liao X, Chen X, Xia Y, Lei T, Sun L, Men W, Chen R, Ma L, Ren X, Wang Y, Wang D, Hu M, Pan Z, Tan S, Gao JH, Qin S, Tao S, Dong Q, He Y (2024) Linking changes in sulcal morphology to cognitive development from childhood to adolescence. 10 December 2024, PREPRINT (Version 1) available at Research Square, doi:10.21203/rs.3.rs-5561682/v1

69. Liang W, Yu Q, Wang W, Dhollander T, Suluba E, Li Z, Xu F, Hu Y, Tang Y, Liu S (2022) A comparative study of the superior longitudinal fasciculus subdivisions between neonates and young adults. Brain Struct Funct 227:2713–2730.

70. Lopez-Persem A, Verhagen L, Amiez C, Petrides M, Sallet J (2019) The human ventromedial prefrontal cortex: sulcal morphology and its influence on functional organization. J Neurosci 39:3627–3639.

71. Luckett PH, Lee JJ, Park KY, Raut RV, Meeker KL, Gordon EM, Snyder AZ, Ances BM, Leuthardt EC, Shimony JS (2023) Resting state network mapping in individuals using deep learning. Front Neurol 13:1055437.

72. Luo AC, Sydnor VJ, Pines A, Larsen B, Alexander-Bloch AF, Cieslak M, Covitz S, Chen AA, Esper NB, Feczko E, Franco AR, Gur RE, Gur RC, Houghton A, Hu F, Keller AS, Kiar G, Mehta K, Salum GA, Tapera T, Xu T, Zhao C, Salo T, Fair DA, Shinohara RT, Milham MP, Satterthwaite TD (2024) Functional connectivity development along the sensorimotor-association axis enhances the cortical hierarchy. Nat Commun 15:3511.

73. Lynch CJ, Elbau IG, Ng T, Ayaz A, Zhu S, Wolk D, Manfredi N, Johnson M, Chang M, Chou J, Summerville I, Ho C, Lueckel M, Bukhari H, Buchanan D, Victoria LW, Solomonov N, Goldwaser E, Moia S, Caballero-Gaudes C, Downar J, Vila-Rodriguez F, Daskalakis ZJ, Blumberger DM, Kay K, Aloysi A, Gordon EM, Bhati MT, Williams N, Power JD, Zebley B, Grosenick L, Gunning FM, Liston C (2024) Frontostriatal salience network expansion in individuals in depression. Nature, 633:624–633.

74. Lyu I, Bao S, Hao L, Yao J, Miller JA, Voorhies W, Taylor WD, Bunge SA, Weiner KS, Landman BA (2021) Labeling lateral prefrontal sulci using spherical data augmentation and context-aware training. NeuroImage 229:117758.

75. Maboudian SA, Willbrand EH, Kelly JP, Jagust WJ, Weiner KS, Alzheimer’s Disease Neuroimaging Initiative (2024) Defining overlooked structures reveals new associations between the cortex and cognition in aging and Alzheimer’s disease. J Neurosci 44:e1714232024.

76. Marek S, Tervo-Clemmens B, Nielsen AN, Wheelock MD, Miller RL, Laumann TO, Earl E, Foran WW, Cordova M, Doyle O, Perrone A, Miranda-Dominguez O, Feczko E, Sturgeon E, Graham A, Hermosillo R, Snider K, Galassi A, Nagel BL, Feldstein Ewing SW, Eggebrecht AT, Garavan H, Dale AM, Greene DJ, Barch DM, Fair DA, Luna B, Dosenbach NUF (2019) Identifying reproducible individual differences in childhood functional brain networks: An ABCD study. Dev Cogn Neurosci 40:100706.

77. Marcus D, Harwell J, Olsen T, Hodge M, Glasser M, Prior F, Jenkinson M, Laumann T, Curtiss S, Van Essen D (2011) Informatics and data mining tools and strategies for the human connectome project. Front Neuroinform 5:4.

78. Markello RD, Misic B (2021) Comparing spatial null models for brain maps. NeuroImage 236:118052.

79. McCarthy P (2022) FSLeyes (1.5.0). doi:10.5281/zenodo.7038115

80. McGugin RW, Newton AT, Tamber-Rosenau B, Tomarken A, Gauthier I (2020) Thickness of deep layers in the fusiform face area predicts face recognition. J Cogn Neurosci 32:1316–1329.

81. McKinney W (2010) Data structures for statistical computing in Python. In *Proceedings of the 9th Python in Science Conference* (eds van der Walt S, Millman J), 51–56.

82. Meredith SM, Whyler NC, Stanfield AC, Chakirova G, Moorhead TW, Job DE, Giles S, McIntosh AM, Johnstone EC, Lawrie SM (2012) Anterior cingulate morphology in people at genetic high-risk of schizophrenia. Eur Psychiatry 27:377–85.

83. Miller JA, Voorhies WI, Lurie DJ, D’Esposito M, Weiner KS (2021) Overlooked tertiary sulci serve as a meso-scale link between microstructural and functional properties of human lateral prefrontal cortex. J Neurosci 41:2229–2244.

84. Miller JA, Weiner KS (2022) Unfolding the evolution of human cognition. Trends Cogn Sci 26:735–737.

85. Moore LA, Hermosillo RJ, Feczko E, Moser J, Koirala S, Allen MC, Buss C, Conan G, Juliano, AC, Marr M, Miranda-Dominguez O, Mooney M, Myers M, Rasmussen J, Rogers CE, Smyser CD, Snider K, Sylvester C, Thomas E, Fair DA, Graham AM (2024) Towards personalized precision functional mapping in infancy. Imaging Neurosci 2:1–20.

86. Natu VS, Arcaro MJ, Barnett MA, Gomez J, Livingstone M, Grill-Spector K, Weiner KS (2021) Sulcal depth in the medial ventral temporal cortex predicts the location of a place-selective region in macaques, children, and adults. Cereb Cortex 31:48–61.

87. Noble S, Spann MN, Tokoglu F, Shen X, Constable RT, Scheinost D (2017) Influences on the test–retest reliability of functional connectivity MRI and its relationship with behavioral utility. Cereb Cortex 27:5415–5429.

88. Parker BJ, Voorhies WI, Jiahui G, Miller JA, Willbrand E, Hallock T, Furl N, Garrido L, Duchaine B, Weiner KS (2023) Hominoid-specific sulcal variability is related to face perception ability. Brain Struct Funct 228:677–685.

89. Parlatini V, Radua J, Dell’Acqua F, Leslie A, Simmons A, Murphy DG, Catani M, Thiebaut de Schotten M (2017) Functional segregation and integration within fronto-parietal networks. NeuroImage 146:367–375.

90. Pedregosa F, Varoquaux G, Gramfort A, Michel V, Thirion B, Grisel O, Blondel M, Prettenhofer P, Weiss R, Dubourg V, Vanderplas J, Passos A, Cournapeau D, Brucher M, Perrot M, Duchesnay E (2011) Scikit-learn: machine learning in Python. J Mach Learn Res 12:2825–2830.

91. Persichetti AS, Shao J, Gotts SJ, Martin A (2024) A functional parcellation of the whole brain in high-functioning individuals with autism spectrum disorder reveals atypical patterns of network organization. Mol Psychiatry, doi:10.1038/s41380-024-02764-6

92. Petrides M (2019) Atlas of the Morphology of the Human Cerebral Cortex on the Average MNI Brain. Academic Press.

93. Power JD, Cohen AL, Nelson SM, Wig GS, Barnes KA, Church JA, Vogel AC, Laumann TO, Miezin FM, Schlaggar BL, Petersen SE (2011) Functional network organization of the human brain. Neuron 72:665–678.

94. Power JD, Barnes KA, Snyder AZ, Schlaggar BL, Petersen SE (2012) Spurious but systematic correlations in functional connectivity MRI networks arise from subject motion. NeuroImage, 59:2142–2154.

95. Power JD, Mitra A, Laumann TO, Snyder AZ, Schlaggar BL, Petersen SE (2014) Methods to detect, characterize, and remove motion artifact in resting state fMRI. NeuroImage 84:320–341.

96. Rajesh A, Seider NA, Newbold DJ, Adeyemo B, Marek S, Greene DJ, Snyder AZ, Shimony JS, Laumann TO, Dosenbach NU, Gordon EM (2024) Structure–function coupling in highly sampled individual brains. Cereb Cortex 34:bhae361.

97. Régis J, Mangin JF, Ochiai T, Frouin V, Riviére D, Cachia A, Tamura M, Samson Y (2005) “Sulcal root” generic model: a hypothesis to overcome the variability of the human cortex folding patterns. Neurol Med Chir (Tokyo) 45:1–17.

98. Reveley C, Seth AK, Pierpaoli C, Silva AC, Yu D, Saunders RC, Leopold DA, Ye FQ (2015) Superficial white matter fiber systems impede detection of long-range cortical connections in diffusion MR tractography. Proc Natl Acad Sci U S A 112:E2820–8.

99. Rosvall M, Bergstrom CT (2008) Maps of random walks on complex networks reveal community structure. Proc Natl Acad Sci 105:1118–1123.

100. Rubinov M, Sporns O (2010) Complex network measures of brain connectivity: Uses and interpretations. NeuroImage 52:1059–1069.

101. Salvo JJ, Holubecki AM, Braga RM (2021) Correspondence between functional connectivity and task-related activity patterns within the individual. Curr Opin Behav Sci 40:178– 188.

102. Sanides F (1964) Structure and function of the human frontal lobe. Neuropsychologia 2:209– 219.

103. Sanides F (1962) Architectonics of the human frontal lobe of the brain. With a demonstration of the principles of its formation as a reflection of phylogenetic differentiation of the cerebral cortex. Monogr Gesamtgeb Neurol Psychiatr 98:1–201.

104. Santacroce F, Cachia A, Fragueiro A, Grande E, Roell M, Baldassarre A, Sestieri C, Committeri G (2024) Human intraparietal sulcal morphology relates to individual differences in language and memory performance. Commun Biol 7:520.

105. Schilling KG, Archer D, Rheault F, Lyu I, Huo Y, Cai LY, Bunge SA, Weiner KS, Gore JC, Anderson AW, Landman BA (2023) Superficial white matter across development, young adulthood, and aging: volume, thickness, and relationship with cortical features. Brain Struct Funct 228:1019–1031.

106. Seabold, S, Perktold J (2010) Statsmodels: Econometric and statistical modeling with Python. In *Proceedings of the 9th Python in Science Conference* (eds van der Walt S, Millman J), 92–96.

107. Seitzman BA, Gratton C, Laumann TO, Gordon EM, Adeyemo B, Dworetsky A, Kraus BT, Gilmore AW, Berg JJ, Ortega M, Nguyen A, Greene DJ, McDermott KB, Nelson SM, Lessov-Schlaggar CN, Schlaggar BL, Dosenbach NUF, Petersen SE (2019) Trait-like variants in human functional brain networks. Proc Natl Acad Sci 116:22851–22861.

108. Shinn M, Hu A, Turner L, Noble S, Preller KH, Ji JL, Moujaes F, Achard S, Scheinost D, Constable RT, Krystal JH, Vollenweider FX, Lee D, Anticevic A, Bullmore ET, Murray JD (2023) Functional brain networks reflect spatial and temporal autocorrelation. Nat Neurosci 26:867–878.

109. Smith DM, Perez DC, Porter A, Dworetsky A, Gratton C (2021) Light through the fog: using precision fMRI data to disentangle the neural substrates of cognitive control. Curr Opin Behav Sci 40:19–26.

110. Smith DM, Kraus BT, Dworetsky A, Gordon EM, Gratton C (2023) Brain hubs defined in the group do not overlap with regions of high inter-individual variability. NeuroImage 277:120195.

111. Stuss DT, Knight RT (2013) Principles of frontal lobe function. Oxford University Press, USA.

112. Sun ZY, Pinel P, Rivière D, Moreno A, Dehaene S, Mangin JF (2016) Linking morphological and functional variability in hand movement and silent reading. Brain Struct Funct 221:3361–3371.

113. Suo X, Ding H, Li X, Zhang Y, Liang M, Zhang Y, Yu C, Qin W (2021) Anatomical and functional coupling between the dorsal and ventral attention networks. NeuroImage 232:117868.

114. Thiebaut de Schotten M, Dell’Acqua F, Forkel SJ, Simmons A, Vergani F, Murphy DGM, Catani M (2011) A lateralized brain network for visuospatial attention. Nat Neurosci 14:1245–1246.

115. Tissier C, Linzarini A, Allaire-Duquette G, Mevel K, Poirel N, Dollfus S, Etard O, Orliac F, Peyrin C, Charron S, Raznahan A, Houdé O, Borst G, Cachia (2018) Sulcal polymorphisms of the IFC and ACC contribute to inhibitory control variability in children and adults. eNeuro 5:197–214.

116. Tomaiuolo F, Giordano F (2016) Cerebal sulci and gyri are intrinsic landmarks for brain navigation in individual subjects: an instrument to assist neurosurgeons in preserving cognitive function in brain tumour surgery (Commentary on Zlatkina et al.). Eur J Neurosci 43:1266–1267.

117. Tomaiuolo F, Raffa G, Morelli A, Rizzo V, Germanó A, Petrides M (2022) Sulci and gyri are topological cerebral landmarks in individual subjects: a study of brain navigation during tumour resection. Eur J Neurosci 55:2037–2046.

118. Tooley UA, Bassett DS, Mackey AP (2022) Functional brain network community structure in childhood: Unfinished territories and fuzzy boundaries. NeuroImage 247:118843.

119. Tustison NJ, Avants BB, Cook PA, Yuanjie Zheng, Egan A, Yushkevich PA, Gee JC (2010) N4ITK: Improved N3 Bias Correction. IEEE Trans Med Imaging 29:1310–1320.

120. Uddin LQ, Supekar KS, Ryali S, Menon V (2011) Dynamic reconfiguration of structural and functional connectivity across core neurocognitive brain networks with development. J Neurosci 31:18578–89.

121. Uddin LQ, Betzel RF, Cohen JR, Damoiseaux JS, De Brigard F, Eickhoff SB, Fornito A, Gratton C, Gordon EM, Laird AR, Larson-Prior L, McIntosh AR, Nickerson LD, Pessoa L, Pinho AL, Poldrack RA, Razi A, Sadaghiani S, Shine JM, Yendiki A, Yeo BTT, Spreng N (2023) Controversies and progress on standardization of large-scale brain network nomenclature. Netw Neurosci 7:864–905.

122. Van Essen DC, Dierker DL (2007) Surface-based and probabilistic atlases of primate cerebral cortex. Neuron 56:209–225.

123. Van Essen DC, Jbabdi S, Sotiropoulos SN, Chen C, Dikranian K, Coalson T, Harwell J, Behrens TE, Glasser MF (2013) Mapping connections in humans and non-human primates: aspirations and challenges for diffusion imaging. In Diffusion MRI, 2nd edition (eds Johansen-Berg H, Behrens TE). Academic Press.

124. Vendetti MS, Bunge SA (2014) Evolutionary and developmental changes in the lateral frontoparietal network: a little goes a long way for higher-level cognition. Neuron 84:906–917.

125. Virtanen, et al. (2020) SciPy 1.0: fundamental algorithms for scientific computing in Python. Nat Methods 17:261–272.

126. Voorhies WI, Miller JA, Yao JK, Bunge SA, Weiner KS (2021) Cognitive insights from tertiary sulci in prefrontal cortex. Nat Commun 12:5122.

127. Wandell BA, Chial S, Backus BT (2000) Visualization and measurement of the cortical surface. J Cogn Neurosci 12:739–752.

128. Waskom ML (2021) seaborn: statistical data visualization. J Open Source Softw 6:3021.

129. Wechsler D (1949) Wechsler Intelligence Scale for Children–Fourth Edition (WISC-IV). The Psychological Corporation.

130. Weiner KS (2023) The hypothesis of fundal cognition. Nat Rev Neurosci 24:521–521.

131. Weiner KS, Golarai G, Caspers J, Chuapoco MR, Mohlberg H, Zilles K, Amunts K, Grill-Spector K (2014) The mid-fusiform sulcus: a landmark identifying both cytoarchitectonic and functional divisions of human ventral temporal cortex. NeuroImage 84:453–465.

132. Weiner KS, Natu VS, Grill-Spector K (2018) On object selectivity and the anatomy of the human fusiform gyrus. NeuroImage 173:604–609.

133. Welker W (1990) Why does cerebral cortex fissure and fold. In Cerebral Cortex (eds Jones EG, Peters A). New York: Plenum Press.

134. Wendelken C, Ferrer E, Ghetti S, Bailey SK, Cutting L, Bunge SA (2017) Frontoparietal structural connectivity in childhood predicts development of functional connectivity and reasoning ability: A large-scale longitudinal investigation. J Neurosci 37:8549– 8558.

135. Wendelken C, O’Hare ED, Whitaker KJ, Ferrer E, Bunge SA (2011) Increased functional selectivity over development in rostrolateral prefrontal cortex. J Neurosci 31:17260– 17268.

136. Willbrand EH, Bunge SA, Weiner KS (2023a) Neuroanatomical and functional dissociations between variably present anterior lateral prefrontal sulci. J Cogn Neurosci 35:1846– 1867.

137. Willbrand EH, Ferrer E, Bunge SA, Weiner KS (2023b) Development of human lateral prefrontal sulcal morphology and its relation to reasoning performance. J Neurosci 43:2552–2567.

138. Willbrand EH, Jackson S, Chen S, Hathaway CB, Voorhies WI, Bunge SA, Weiner KS (2024) Cognitive relevance of an evolutionarily new and variable prefrontal structure. Brain Struct Funct 229:387–402.

139. Willbrand EH, Tsai Y-H, Gagnant T, Weiner KS (2023c) Updating the sulcal landscape of the human lateral parieto-occipital junction provides anatomical, functional, and cognitive insights. eLife 12:RP90451.

140. Willbrand EH, Voorhies WI, Yao JK, Weiner KS, Bunge SA (2022) Presence or absence of a prefrontal sulcus is linked to reasoning performance during child development. Brain Struct Funct 227:2543–2551.

141. Woolgar A, Parr A, Cusack R, Thompson R, Nimmo-Smith I, Torralva T, Roca M, Antoun N, Manes F, Duncan J (2010) Fluid intelligence loss linked to restricted regions of damage within frontal and parietal cortex. Proc Natl Acad Sci 107:14899–14902.

142. Yao JK, Voorhies WI, Miller JA, Bunge SA, Weiner KS (2023) Sulcal depth in prefrontal cortex: A novel predictor of working memory performance. Cereb Cortex 33:1799– 1813.

143. Yeo BTT, Krienen FM, Sepulcre J, Sabuncu MR, Lashkari D, Hollinshead M, Roffman JL, Smoller JW, Zöllei L, Polimeni JR, Fischl B, Liu H, Buckner RL (2011) The organization of the human cerebral cortex estimated by intrinsic functional connectivity. J Neurophysiol 106:1125–1165.

144. Zhang Y, Brady M, Smith S (2001) Segmentation of brain MR images through a hidden Markov random field model and the expectation-maximization algorithm. IEEE Trans Med Imaging 20:45–57.

145. Zhi D, King M, Hernandez-Castillo CR, Diedrichsen J (2022) Evaluating brain parcellations using the distance-controlled boundary coefficient. Hum Brain Mapp 43:3706–3720.

146. Zilles K, Palomero-Gallagher N, Amunts K (2013) Development of cortical folding during evolution and ontogeny. Trends Neurosci 36:275–284.

147. Zilles K, Schleicher A, Langemann C, Amunts K, Morosan P, Palomero-Gallagher N, Schormann T, Mohlberg H, Bürgel U, Steinmetz H, Schlaug G (1997) Quantitative analysis of sulci in the human cerebral cortex: development, regional heterogeneity, gender difference, asymmetry, intersubject variability and cortical architecture. Human Brain Mapp 5:218–221.

148. Zlatkina V, Petrides M (2014) Morphological patterns of the intraparietal sulcus and the anterior intermediate parietal sulcus of Jensen in the human brain. Proc R Soc B Biol Sci 281:20141493.

149. Zlatkina V, Amiez C, Petrides M (2016) The postcentral sulcal complex and the transverse postcentral sulcus and their relation to sensorimotor functional organization. Eur J Neurosci 43:1268–1283.

